# High-speed single-molecule imaging reveals signal transduction by induced transbilayer raft phases

**DOI:** 10.1101/2020.06.20.162545

**Authors:** Ikuko Koyama-Honda, Takahiro K. Fujiwara, Rinshi S. Kasai, Kenichi G. N. Suzuki, Eriko Kajikawa, Hisae Tsuboi, Taka A. Tsunoyama, Akihiro Kusumi

## Abstract

Using single-molecule imaging with enhanced time resolutions down to 5 ms, we found that CD59-cluster rafts and GM1-cluster rafts stably induced in the outer leaflet of the plasma membrane (PM), which triggered the activation of Lyn, H-Ras, and ERK, continually recruited Lyn and H-Ras right beneath them in the inner leaflet, with dwell lifetimes of <0.1 s. The detection was possible due to the enhanced time resolutions employed here. The recruitment depended on the PM cholesterol and saturated alkyl chains of Lyn and H-Ras, whereas it was blocked by the non-raftophilic transmembrane protein moiety and unsaturated alkyl chains linked to the inner-leaflet molecules. Since GM1-cluster rafts recruited Lyn and H-Ras as efficiently as CD59-cluster rafts, and the protein moieties of Lyn and H-Ras were not required for the recruitment, we conclude that the transbilayer raft phases induced by the outer-leaflet stabilized rafts recruit lipid-anchored signaling molecules by lateral raft-lipid interactions, and thus serve as a key signal transduction platform.

**Summary:** High-speed single-molecule imaging indicated that CD59-cluster rafts and GM1-cluster rafts stably induced in the plasma membrane outer leaflet generated nano-scale transbilayer raft phases, which continually and transiently recruited Lyn and H-Ras in the inner leaflet by cooperative raft-lipid interactions.

## INTRODUCTION

In the human genome, more than 150 protein species have been identified as glycosylphosphatidylinositol (GPI)-anchored proteins, in which the protein moieties located at the extracellular surface of the plasma membrane (PM) are anchored to the PM by way of GPI, a phospholipid (Kinoshita and Fujita, 2016). Many GPI-anchored proteins are receptors, and thus referred to as GPI-anchored receptors (GPI-ARs). A GPI-anchored structure appears paradoxical for receptors because it spans only halfway through the membrane, and yet, to function as a receptor, it has to relay the signal from the outside environment to the inside of the cell (**Fig. 1A**). “Raft domains”, the PM domains on the space scales from a few nanometers up to several hundred nanometers that are built by cooperative interactions of cholesterol and molecules with saturated alkyl chains of C16 or longer, as well as by their exclusion from the bulk unsaturated-chain-enriched domains (Kusumi et al., 2020; Levental et al., 2020), have been implied in the signaling process of GPI-ARs across the plasma membrane (PM; Omidvar et al., 2006; Suzuki et al., 2007b; Paulick and Bertozzi, 2008; Eisenberg et al., 2011; Fessler and Parks, 2011; Lingwood et al., 2011; Suzuki et al., 2012; Kusumi et al., 2014; Raghupathy et al., 2015). Nevertheless, exactly how raft domains or raft-based lipid interactions participate in the transbilayer signal transduction of GPI-ARs remains unknown. Indeed, raft-based interactions might even be involved in the signal transduction by transmembrane receptors (Coskun et al., 2011; Chung et al., 2016; Shelby et al., 2016).

**Figure 1.**
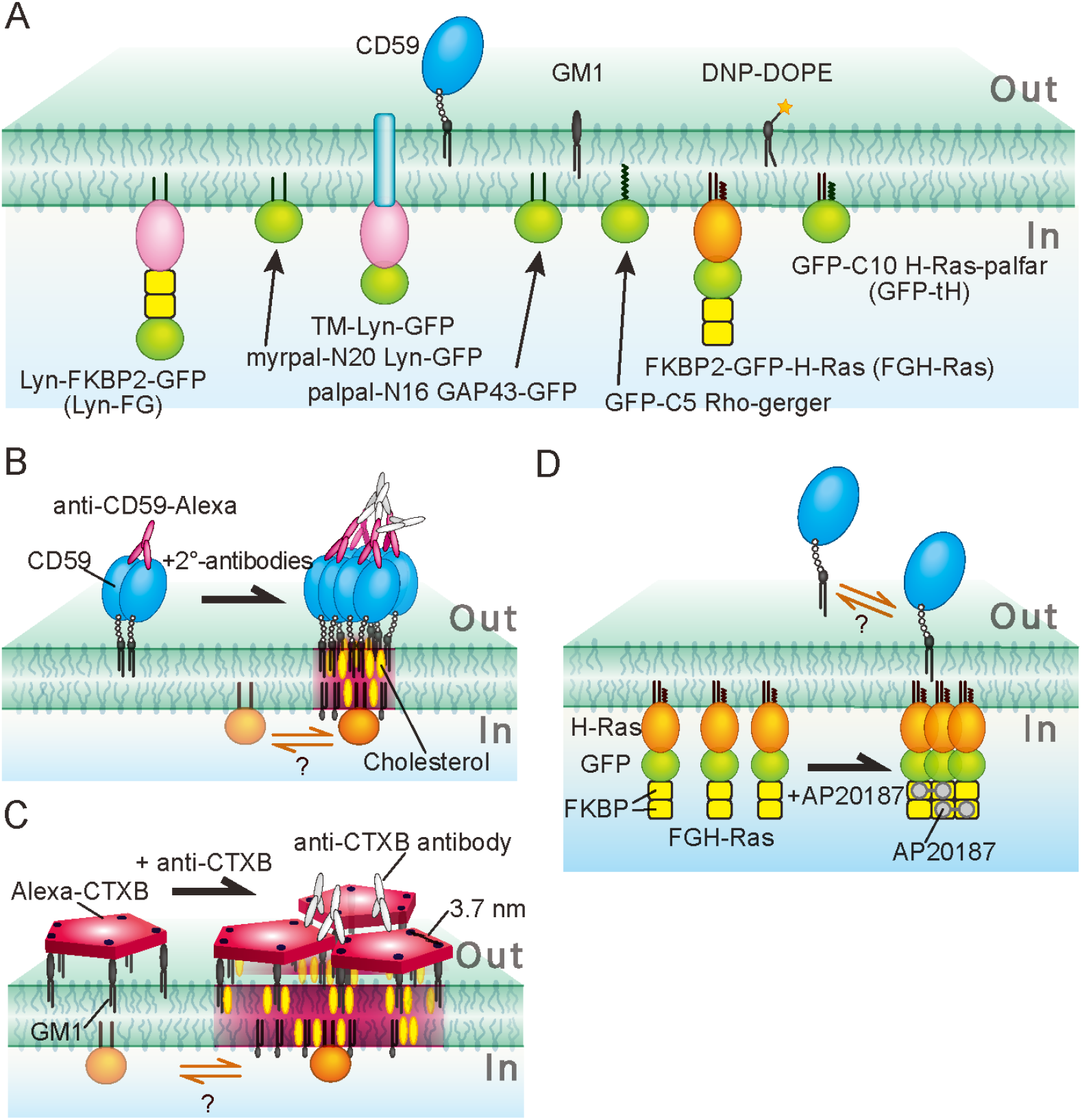
Outer- and inner-leaflet lipid-anchored molecules employed in this study and their crosslinking schemes. **(A)** The outer-leaflet molecules employed in this work were a prototypical GPI-AR, CD59, a prototypical ganglioside, GM1, and a prototypical non-raft phospholipid, L-α-dioleoylphosphatidylethanolamine (DOPE) conjugated by DNP (DNP-DOPE). The inner-leaflet molecules examined here were (G and GFP represent EGFP) the following. Lyn-FKBP2-GFP (Lyn-FG): Lyn conjugated, at its C-terminus, to two molecules of FK506 binding protein (FKBP) in series and then to GFP. Myrpal-N20Lyn-GFP: myristoyl, palmitoyl-anchored-Lyn peptide conjugated to GFP, where the peptide was the 20 amino-acid N-terminal sequence of Lyn, which contains the conjugation sites for both myristoyl and palmitoyl chains. Transmembrane (TM)-Lyn-GFP: the TM mutant of Lyn-GFP, in which the TM domain of a prototypical non-raft molecule LDL-receptor was conjugated to the N-terminus of the full length Lyn-GFP (which cannot be fatty-acylated). Palpal-N16GAP43-GFP: palmitoyl, palmitoyl-anchored GAP43 peptide conjugated to GFP, in which the peptide was the 16 amino-acid N-terminal sequence of GAP43 containing two palmitoylation sites (likely to be raft-associated). GFP-C5Rho-geranylgeranyl: GFP anchored by a geranylgeranyl chain, in which GFP was conjugated at its C-terminus to the 5 amino-acid C-terminal sequence of Rho, which contains a site for attaching an unsaturated geranylgeranyl chain (likely to be *non*-raft-associated). FKBP2-GFP-H-Ras (FGH-Ras): H-Ras chimera molecule, in which two tandem FKBP molecules linked to GFP were then conjugated to H-Ras. GFP-C10H-Ras-palfar (GFP-tH): GFP linked to the 10 amino-acid C-terminal sequence of H-Ras, containing two sites for palmitoylation and a site for farnesylation. These molecules were expressed and observed in live HeLa cells. **(B, C, D)** The schemes for clustering (crosslinking) CD59 **(B)**, GM1 (**C**), and FGH-Ras (D). CD59 was clustered by the sequential additions of anti-CD59 monoclonal antibody IgG labeled with the fluorescent dye Alexa633 (A633) and secondary antibodies (**B**). GM1 was clustered by the sequential additions of CTXB conjugated with A633 and anti-CTXB antibodies (**C**). FGH-Ras (as well as Lyn-FG) was clustered by the addition of AP20187 (crosslinker for FKBP) (**D**). After the induction of clustering of these molecules, the possible recruitment of lipid-anchored molecules in the other leaflet of the PM at these clusters was examined.

In giant unilamellar vesicles undergoing liquid-ordered (Lo)/liquid-disordered (Ld)-phase separation, the Lo/Ld-phase domains in the outer leaflet spatially match the same domains in the inner leaflet, indicating strong interbilayer coupling due to phase separation across the bilayer (Collins and Keller, 2008; Blosser et al., 2015). In living cells, the long-chain phosphatidylserine present in the PM inner leaflet was proposed to play key roles in the transbilayer coupling (Raghupathy et al., 2015). However, the mechanisms of transbilayer coupling in the PM for the induction of signal transduction are not well understood.

Using CD59 as a prototypical GPI-AR, our previous single fluorescent-molecule imaging showed that nanoparticle-induced CD59 clusters form stabilized raft domains with diameters on the order of 10 nm in the PM outer leaflet, which in turn continually recruit Giα, Lyn, and PLCγ2 molecules one after another in a manner dependent on raft-lipid interactions, triggering the IP_3_-Ca^2+^ signaling pathway. Therefore, the CD59 clusters were termed “CD59-cluster signaling rafts” or simply “CD59-cluster rafts” (Stefanova et al., 1991; Suzuki et al., 2007a; Suzuki et al., 2007b; Simons and Gerl, 2010; Suzuki et al., 2012; Zurzolo and Simons, 2016). Importantly, the recruitment of cytoplasmic signaling molecules at the CD59 signaling rafts occurred transiently, in the time scale on the order of fractions of a second (in the following, we will use the expression “recruitment of signaling molecules ‘at’ CD59 clusters” rather than “the recruitment ‘to’ CD59 clusters”, because our imaging method could not directly show the binding of the signaling molecules located in the inner leaflet to the CD59 clusters located in the outer leaflet). Furthermore, our single-molecule study revealed that, although gangliosides and sphingomyelins are always present in the CD59-cluster signaling rafts, each lipid molecule associates with the CD59-cluster raft for only 50 ~ 100 ms (Komura et al., 2016; Kinoshita et al., 2017).

Meanwhile, the time resolution of the single-molecule imaging method used to detect such transient colocalization events was only 33 ms. In the present study, we greatly enhanced the imaging time resolutions down to 5.0 and 6.45 ms, an improvement by factors of 6.7 and 5.2, respectively, and thus substantially refined the detection of cytoplasmic signaling molecule colocalizations with CD59-cluster rafts (and GM1-cluster rafts). To the best of our knowledge, these are likely to be the fastest simultaneous, two-color, single-molecule observations ever performed. We previously found Lyn recruitment at CD59-cluster rafts, but in the present research, by applying single-molecule imaging at enhanced time resolutions and using various lipid-anchored cytoplasmic molecules, including Lyn, H-Ras, and four artificial, designed molecules, as well as by employing the stabilized ganglioside GM1-cluster rafts, in addition to the CD59-cluster rafts, we sought to unravel the mechanisms by which cytoplasmic lipid-anchored signaling molecules in the PM inner leaflet are recruited at CD59-cluster rafts and GM1-cluster rafts formed in the PM outer leaflet.

In addition to the well-known function of CD59 to protect normal cells in the body against self-attack by the membrane-attack complement complex (MACC), CD59 is involved in tumor growth. First, CD59 renders autologous carcinoma cells insensitive to the MACC action, providing tumor cells with a key strategy to evade the immune system (Morgan et al., 1998; Carter and Lieber, 2014). Second, the MACC-induced CD59 clusters activate the extracellular signal-regulated kinase (ERK) signaling pathway, thus enhancing tumor cell proliferation (Jurianz et al., 1999). Therefore, the basic understanding of CD59 signaling, particularly the Lyn (Src-family kinase) signaling to trigger the IP3-Ca^2+^ pathway for protection against MACC binding, as well as the signaling cascades for ERK activation by way of Lyn and Ras (Cantrell, 1996; Bertotti et al., 2006; Harita et al., 2008; Wang et al., 2011; Suzuki et al., 2012; Croucher et al., 2013; Dorard et al., 2017), would be useful for developing methods to regulate CD59 function, eventually leading to better therapeutic outcomes in oncology by suppressing ERK activities and reversing complement resistance (Carter and Lieber, 2014).

In the present research, we first aimed to unravel how the CD59-cluster rafts in the PM outer leaflet recruit the downstream intracellular lipid-anchored signaling molecules Lyn and H-Ras, located in the PM inner leaflet. Lyn is anchored to the PM inner leaflet by a myristoyl chain and a palmitoyl chain, whereas H-Ras is anchored by two palmitoyl chains and a farnesyl chain (**Fig. 1A**). Since CD59 cannot directly interact with and activate Lyn and H-Ras, and since Lyn and H-Ras are proposed to be raft-domain-associated in the PM inner leaflet (Field et al., 1997; Sheets et al., 1999; Prior et al., 2001; Prior et al., 2003), we paid special attention to raft-lipid interactions as a recruiting mechanism (Wang et al., 2005), while also considering protein-protein interactions (Douglass and Vale, 2005) (**Fig. 1B**).

Second, to directly examine the possibility that the signal transfer from the PM outer leaflet to the inner leaflet is mediated by raft-lipid interactions, we crosslinked the prototypical raft lipid ganglioside GM1 in the outer leaflet, to examine whether GM1 clusters could recruit Lyn and H-Ras in the inner leaflet (**Fig. 1A, C**). Many studies have examined the cytoplasmic signals triggered by GPI-AR stimulation and GM1 clustering in raft-dependent manners (Pyenta et al., 2001; McKerracher and Winton, 2002; Wang et al., 2005; Todeschini et al., 2008; Fujita et al., 2009; Um and Ko, 2017), although the results varied considerably. In contrast, very few studies have investigated the actual recruitment of cytoplasmic lipid-anchored signaling molecules at the stabilized nano-raft domains formed in the PM outer leaflet (Harder et al., 1998; Suzuki et al., 2007a; Suzuki et al., 2007b; Suzuki et al., 2012), and particularly the molecular dynamics of the recruitment in live cells. In the present study, as a control, we induced the clustering of lipid-anchored Lyn or H-Ras in the PM inner leaflet, and observed whether this could induce the recruitment of CD59 and GM1 in the PM outer leaflet (**Fig. 1A, D**).

## RESULTS

### Antibody-induced CD59 clusters in the PM and their ERK activation

First, we improved the time resolution of our home-built single-molecule imaging station, described previously (Koyama-Honda et al., 2005; Komura et al., 2016; Kinoshita et al., 2017). The improvements were accomplished by employing two kinds of camera systems that can operate at higher frame rates (**Materials and methods**) and modifying the single-molecule imaging station, by using lasers with higher outputs and tuning the excitation optics. As a result, the time resolution was enhanced from 33.3 ms (30 Hz) to 5.0 or 6.45 ms (200 or 155 Hz, respectively, which is faster than normal video rate by factors of 6.7 and 5.2, respectively), with frame sizes of 640 x 160 pixels and 653 x 75 pixels, respectively. We employed the same two cameras for performing simultaneous, two-color, single-molecule imaging (**Materials and methods**). Throughout this work, all of the microscopic observations of CD59-cluster rafts (A633-tagged) and the downstream cytoplasmic signaling molecules (fused to EGFP, which is simply called GFP for conciseness) were performed simultaneously in the bottom (basal) PM of HeLa cells.

CD59-cluster signaling rafts were formed by the addition of the primary (anti-CD59 monoclonal antibody IgG [mAb] conjugated with Alexa633 [A633]) and secondary antibodies, according to previous reports (Field et al., 1997; Janes et al., 1999; Chen and Williams, 2013). Using this method, CD59 clusters could be formed in both the apical and basal PMs, whereas in our previous method of using nano-particles to induce CD59 clusters, due to the non-accessibility of the particles in the space between the basal PM and the coverslip, CD59 clusters were formed only in the *apical*PM. Therefore, in this study, we observed the CD59 clusters and signaling molecules in the *basal*PM, which enabled observations with improved signal-to-noise ratios. To better observe the short-term colocalizations of lipid-anchored signaling molecules with CD59-cluster rafts, we hoped to slow down the colocalization processes, and therefore all microscopic observations were performed at 27°C. These observations were conducted within 10 min after the addition of the secondary antibodies, when more than 92% of the CD59 clusters were located outside caveolae (**Fig. S1A**).

The CD59 clusters consisted of an average of ~10 molecules of CD59 (**Fig. 2A, B**; **Materials and methods**). Since CD59 is anchored to the PM outer leaflet by way of two saturated, long alkyl chains, the CD59 clusters employed here would contain an average of 20 saturated long alkyl chains of CD59 in the small cross-sectional area of the CD59 cluster. The CD59 clusters diffused at a three-fold slower rate than monomeric CD59 (labeled with anti-CF59-Fab-A633; **Fig. 2C, D**). Since we used the dye (A633)-conjugated antibody (and the secondary antibodies) to induce CD59 clusters, the recording periods were quite limited due to photobleaching (~0.51 s) and signal-to-noise ratios for observing the CD59 clusters were worse as compared with our previous observations using fluorescent nano-particles, in the present study we could not detect stimulation-induced temporary arrest of lateral diffusion (STALL), and the CD59 clusters appeared to simply undergo slow diffusion.

**Figure 2.**
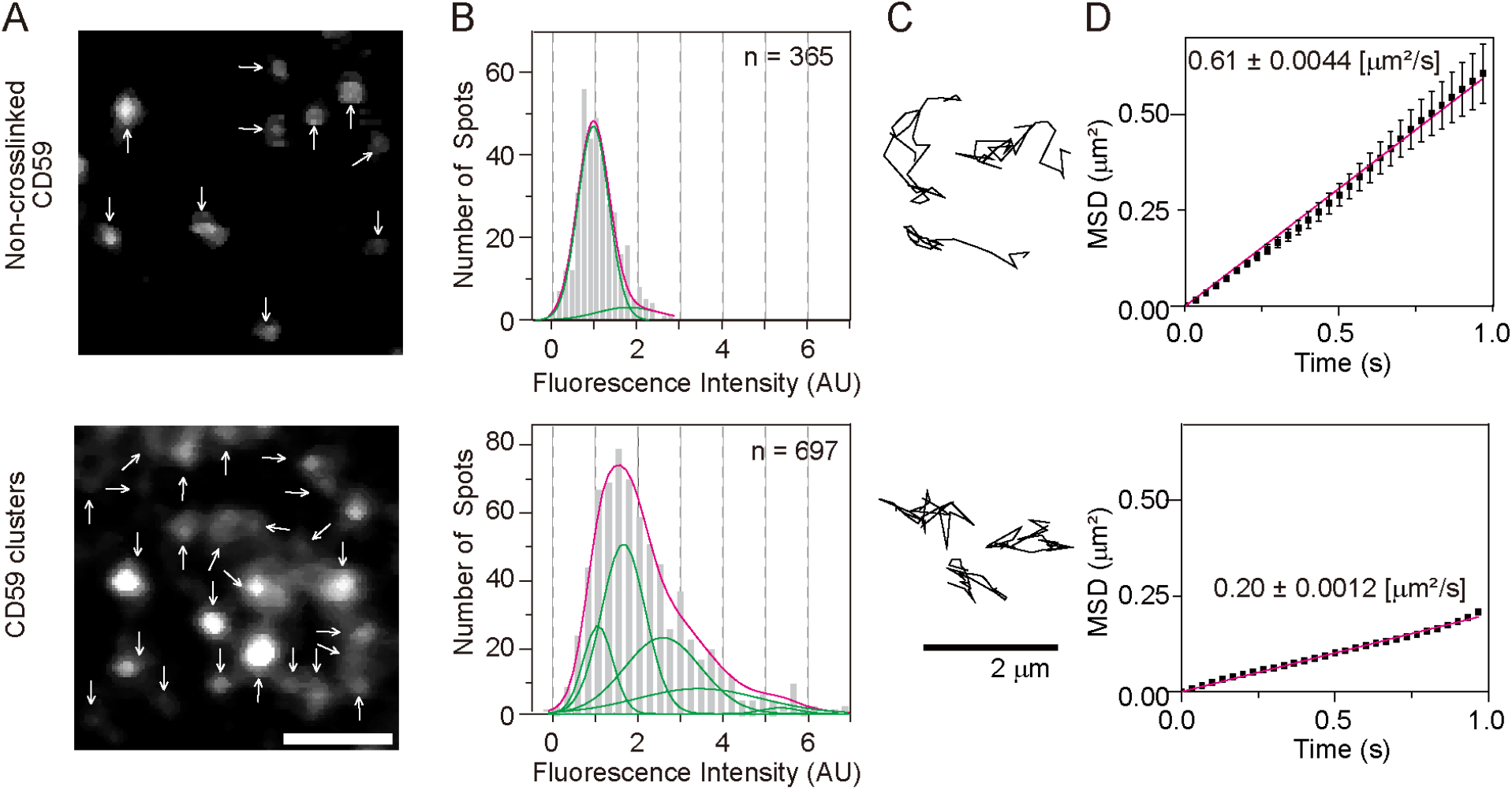
CD59 clusters in the PM outer leaflet contained an average of ~10 CD59 molecules and diffused slowly. **(A)** Fluorescence images of non-crosslinked CD59 bound by A633-anti-CD59 Fab (dye-to-protein mole ratio = 0.27; top) and CD59 clusters induced by the sequential additions of A633-anti-CD59 IgG (a dye-to-protein mole ratio = 0.63) and the secondary antibodies (bottom), obtained at single-molecule sensitivities. Arrows indicate all of the detected fluorescent spots in each image. **(B)** Histograms showing the distributions of the signal intensities of individual fluorescent spots of non-crosslinked CD59 (Fab-A633 probe; top) and CD59 clusters (bottom). Based on these histograms, we concluded that each CD59 cluster contained an average of ~10 CD59 molecules (see **Materials and methods**), although the number distributions would be quite broad. **(C)** Typical trajectories of non-crosslinked CD59 (top) and CD59 clusters (bottom) for 0.2 s, obtained at a time resolution of 6.45 ms. **(D)** Ensemble-averaged mean-square displacements (MSDs) plotted against time, suggesting that in the time scale of 1 s, both non-crosslinked and clustered CD59 undergo effective simple-Brownian diffusion, and the diffusion is slowed by a factor of about three after antibody-induced clustering.

CD59 clustering triggered the signaling cascade to activate the ERK 1/2 kinases (**Fig. 3**), in agreement with the previous finding (Jurianz et al., 1999). The signaling pathways leading to ERK activation could involve the small G-protein H-Ras, as well as Lyn (Bertotti et al., 2006; Harita et al., 2008; Porat-Shliom et al., 2008; Wang et al., 2011; Croucher et al., 2013; Dorard et al., 2017). Therefore, we performed direct single-molecule observations of the recruitment of both Lyn kinase and H-Ras to the CD59-cluster signaling rafts. We had previously detected Lyn recruitment at CD59 clusters (Suzuki et al., 2007a; Suzuki et al., 2007b), but in the present research we focused on understanding the recruitment mechanism by using other related molecules and H-Ras, as well as by employing single-molecule observations with improved time resolutions.

**Figure 3.**
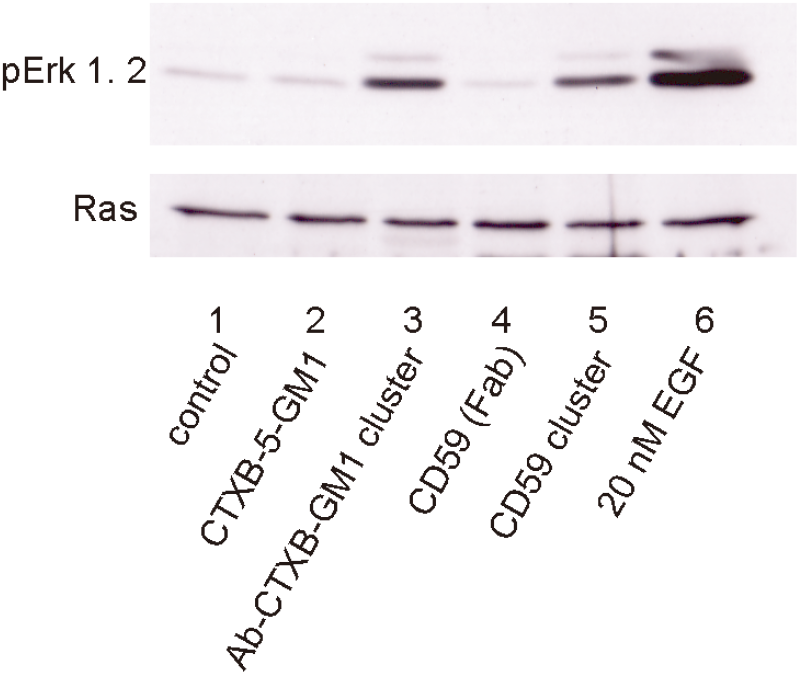
Both CD59 clusters and GM1 clusters induced by the sequential additions of CTXB and its polyclonal antibodies (Ab-CTXB-GM1 clusters) induced Erk phosphorylation (activation). Note that the simple clustering of five GM1 molecules by CTXB (CTXB-5-GM1) failed to trigger ERK activation. Western blotting was performed by using anti-phosphorylated Erk antibodies (top), with anti-H-Ras antibodies as the loading controls (bottom). The addition of 20 nM EGF was used as a positive control for Erk activation.

### Lyn is continually and transiently recruited at CD59-cluster rafts one molecule after another, but not at non-clustered CD59

Lyn is anchored to the PM inner leaflet by myristoyl and palmitoyl chains conjugated to its N-terminus. The Lyn-FG used here for single-molecule observations would be functional, as it could be phosphorylated in RBL-2H3 cells after antigen (DNP) stimulation (**Fig. S2A**). Virtually all of the Lyn-FG molecules (**Fig. 1A**) on the PM inner leaflet underwent thermal diffusion, with a mean diffusion coefficient (in the time scale of 124 ms) of 0.76 ± 0.0019 μm^2^/s (**Fig. 4A**).

**Figure 4.**
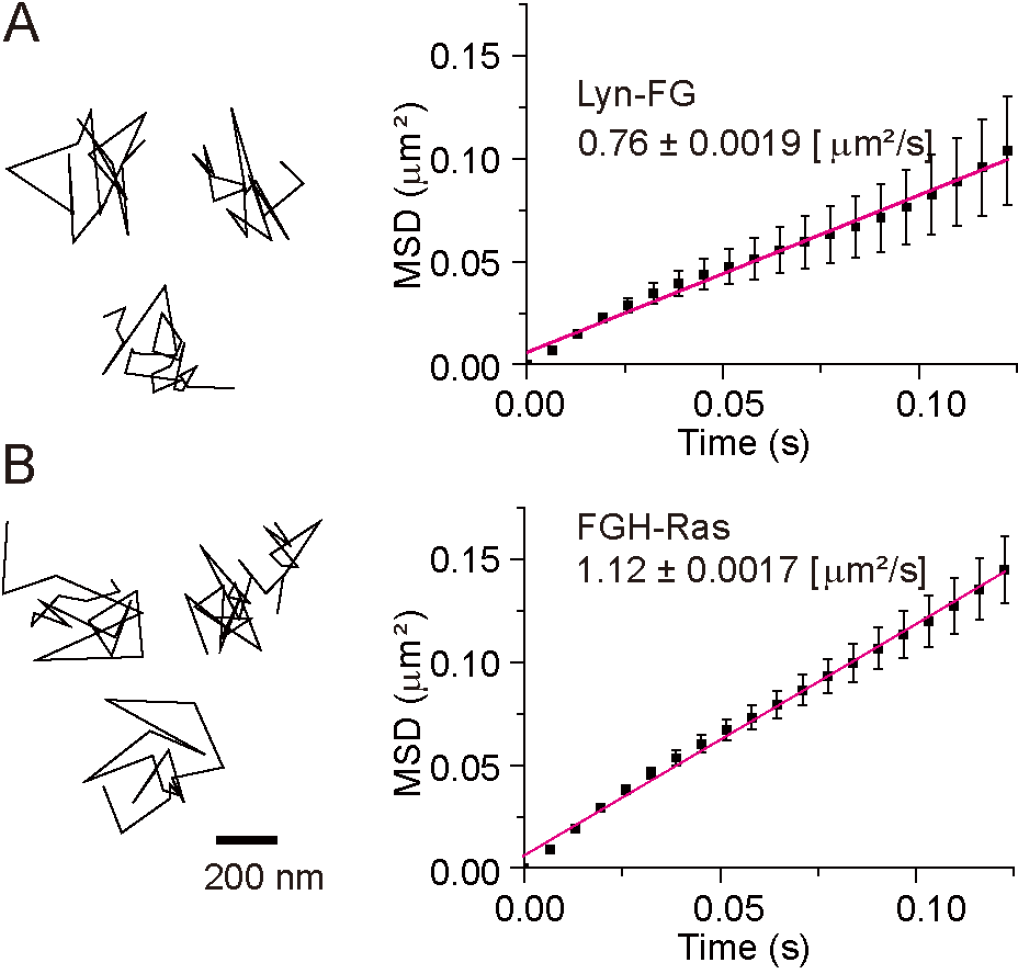
Lyn-FG and FGH-Ras molecules underwent simple-Brownian diffusion in/on the inner PM leaflet as observed at a 6.45-ms resolution, when they were not colocalized with CD59 clusters or Ab-CTXB-GM1 clusters. Representative trajectories of single Lyn-FG and FGH-Ras molecules and the ensemble-averaged MSDs plotted against Δ*t* for Lyn-FG and FGH-Ras (**A** and **B** based on 109 and 456 trajectories, respectively). Their mean diffusion coefficients are shown in the figure.

Simultaneous two-color single-molecule observations revealed that Lyn-FG molecules diffusing in the inner leaflet were continually recruited at CD59 clusters located in the PM outer leaflet, one molecule after another. Importantly, the dwell time of each Lyn-FG molecule at the CD59 cluster was on the order of 0.1 s (**Fig. 5; Video 1**). Quantitative detection of colocalizations was performed by using our previously developed definition, in which fluorescent spots with two different colors are located within 150 nm (Koyama-Honda et al., 2005). Although the colocalization distance of 150 nm is clearly much greater than the sizes of the interacting molecules, which would generally be on the order of several nanometers, the colocalization analysis is still useful for detecting molecular interactions for the following reason. Unassociated molecules may track together by chance over short periods of time for short distances, but the probability of this occurring for multiple frames is small. Therefore, longer colocalization durations imply the presence of molecular interactions between the two molecules, rather than incidental encounters (although molecular interactions are initiated by incidental encounters; see **Materials and methods**).

**Figure 5.**
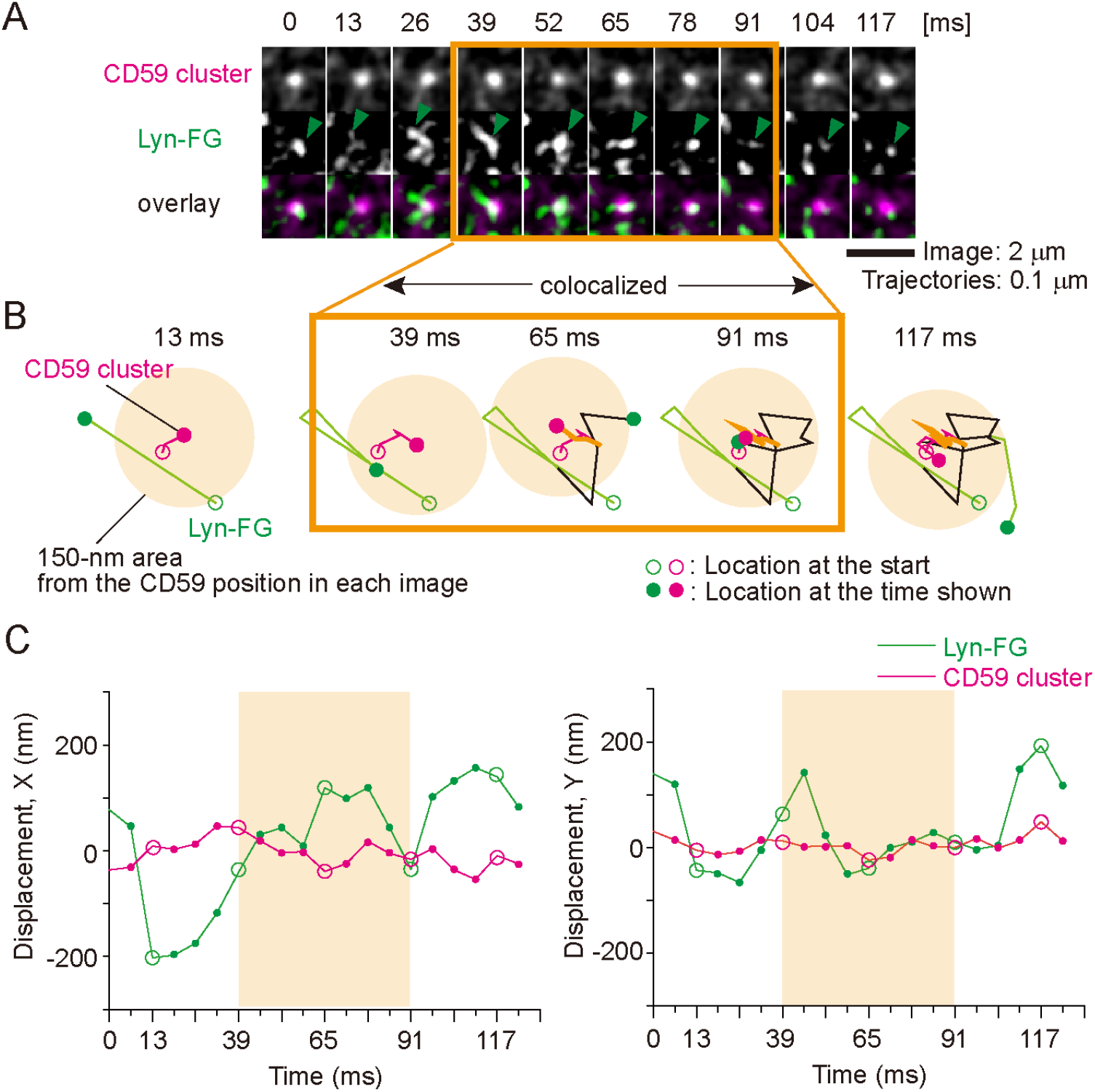
High-speed, simultaneous two-color, single-molecule imaging showed transient recruitment of Lyn-FG in/on the inner leaflet at CD59 clusters located in/on the outer leaflet. **(A)** Typical single-molecule image sequences (6.45-ms resolution; every other image is shown), showing the colocalization of a CD59 cluster (top row and magenta spots in the bottom row) and a single molecule of Lyn-FG (green arrowheads in the middle row and green spots in the bottom row). Lyn-FG spots appear brighter during colocalization due to slower diffusion. **(B)** The trajectories of the CD59 cluster (magenta) and the Lyn-FG molecule (green) shown in **(A)**. These molecules became colocalized (the orange circular region with a radius of 150 nm around the CD59 cluster position) between 39 and 91 ms (52 ms, orange box). **(C)** Another display of the colocalization event shown in **(A)** and **(B)**, showing the displacements of a Lyn-FG molecule and a CD59 cluster along the x and y axes (left and right, respectively) from the average position of the CD59 cluster during the colocalization period, plotted against time. Circles indicate the times employed in **(B)**.

Each time we detected a colocalization event of a Lyn-FG molecule with a CD59 cluster, we measured its duration, and after observing sufficient numbers of colocalization events, we obtained a histogram showing the distribution of colocalized durations for Lyn-FG and CD59 clusters (**Fig. 6A-a; Materials and methods**). However, this duration histogram must also contain the colocalization events due to incidental close encounters of molecules within 150 nm, without any molecular interactions. To obtain the histogram of incidental colocalization durations, the image obtained in the longer-wavelength channel (A633) was shifted toward the right by 20 pixels (1.0 and 1.19 μm, depending on the camera) and then overlaid on the image obtained in the GFP channel (“shifted overlay”). The duration histogram for incidental colocalization, called *h*(incidental-by-shift), could effectively be fitted with a single exponential function with a decay time constant *τ*_1_ of 15 ± 0.93 ms (throughout this report, the SEM of the dwell lifetime is provided by the fitting error of the 68.3% confidence limit for the decay time constant).

**Figure 6.**
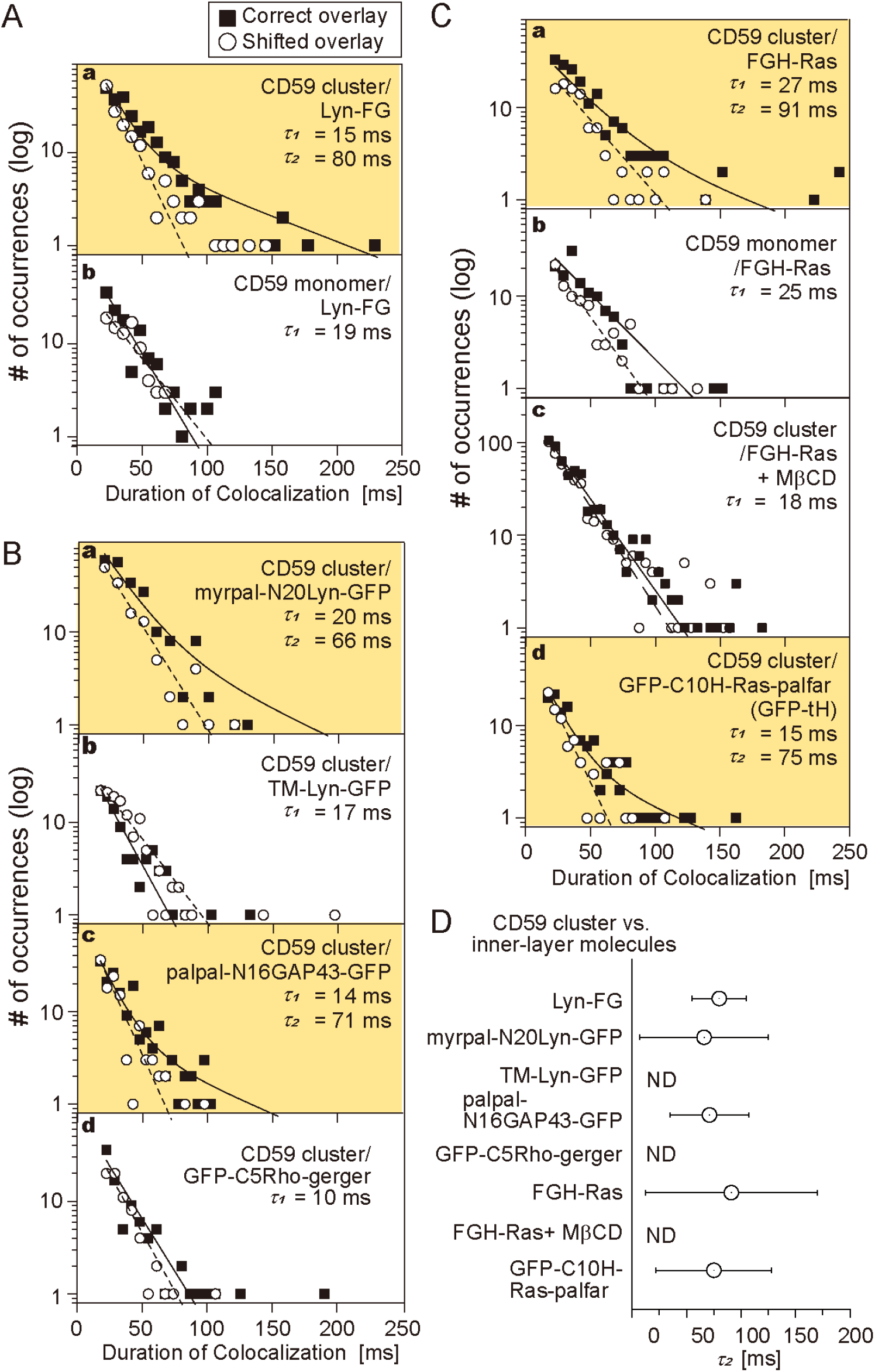
Lyn-FG, FGH-Ras, and other lipid-anchored raftophilic molecules were recruited at CD59 clusters, but not at non-crosslinked CD59. The distributions (histograms) of the colocalization durations for the “correct” and “shifted” overlays, shown in semi-log plots. The histograms for shifted overlays were fitted by a single exponential function (dashed line), and those for the correct overlays were fitted by the sum of two exponential functions (solid line), with the shorter time constant set to *τ*_1_ obtained from the histogram of the shifted overlay. The boxes highlighted in orange contain histograms that could be better fitted with the sum of two exponential decay functions, rather than a single exponential function. The values of *τ*_1_ and *τ*_2_ are indicated in each box. See **Supplementary Table 1** for statistical parameters. **(A)** Lyn-FG was recruited at CD59 clusters, but not at non-crosslinked CD59. **(B)** Recruitment of Lyn-related molecules and other lipid-anchored cytoplasmic model proteins at CD59 clusters: myrpal-N20LynGFP and palpal-N16GAP43-GFP were recruited, but TM-Lyn-GFP and GFP-C5Rho-gerger were not. **(C)** FGH-Ras was recruited at CD59 clusters, but not at non-crosslinked CD59 (**a, b**), and FGH-Ras recruitment at CD59 clusters depended on the PM cholesterol (**c**). Meanwhile, GFP-C10H-Ras-palfar [GFP-tH]) was recruited at CD59 clusters. **(D)** Summary of the bound lifetimes (*τ*_2_’s) of Lyn-FG, FGH-Ras, and other cytoplasmic lipid-anchored signaling molecules at CD59 clusters. The differences in *τ*_2_ values are non-significant. ND, not detected.

The distribution of the durations obtained for correctly overlaying the Lyn-FG movies and CD59-cluster movies was significantly different from that for the shifted overlay *(P* = 0.00076 using the Brunner-Munzel test; Brunner and Munzel, 2000; throughout this report, the Brunner-Munzel test was used for the statistical analysis, and all statistical parameters are summarized in **Supplementary Tables 1-3**). The histogram of colocalization durations for Lyn-FG at CD59 clusters could be fitted with the sum of two exponential functions with decay time constants of *τ*_1_ and *τ*_2_. In the fitting, *τ*_1_ was preset as the decay time constant determined from *h*(incidental-by-shift) and *τ*_2_ was determined as a free parameter. In the previous studies employing normal video rate (30 Hz; 33-ms resolution), due to insufficient time resolutions, such distinct components could not be observed in the colocalization duration histogram.

As described in **Materials and methods**, *τ*_2_ directly represents the binding duration (inverse off-rate assuming simple zero-order dissociation kinetics for Lyn-FG from the CD59 cluster). However, note that, in the scope of this report, we paid more attention to the presence of two components in the colocalization duration histograms, rather than the actual values of *τ*_2_.

In the case of the colocalizations of Lyn-FG with CD59-cluster rafts, the fitting provided a *τ*_2_ of 80 ± 25 ms (**Supplementary Table 1**). Both *τ*_1_ and *τ*_2_ for all of the molecules investigated here were much shorter than the photobleaching lifetimes of GFP and A633 (>400 ms; i.e., 62 or 80 image frames), and therefore no corrections for photobleaching were performed in this research. The 80-ms dwell lifetime of Lyn at CD59 clusters is shorter than that observed previously (median = 200 ms; Suzuki et al., 2007a), probably due to the improved time resolutions and signal-to-noise ratios (previously, shorter colocalizations were likely missed), as well as the different ways of forming CD59 clusters. Therefore, this result indicates that Lyn is recruited at CD59-cluster rafts more transiently than we previously evaluated.

Next, the colocalizations of Lyn-FG with non-clustered CD59 (labeled with anti-CD59 Fab-A633) were examined. The duration histogram obtained by the correct overlay was almost the same as *h*(incidental-by-shift) (*P* = 0.86; *τ*_1_ = 19 ms; **Fig. 6A-b** and **Supplementary Table 1**), and it was significantly different from the histogram with CD59 clusters (*P* = 0.018).

### Lyn recruitment at CD59 clusters requires raft-lipid interactions

Next, we asked whether raft-lipid interactions and protein-protein interactions are required for recruiting Lyn-FG at CD59 clusters. First, we examined the recruitment of myrpal-N20(Lyn)-GFP (**Fig. 1A**), which was proposed to be associated with raft domains (Pyenta et al., 2001). The duration histogram for the colocalizations of myrpal-N20(Lyn)-GFP molecules with CD59 clusters clearly exhibited two components (significant difference from *h*(incidental-by-shift), *P* = 0.025), with a statistically non-significant (*P* < 0.068) 18% reduction in *τ*_2_ (**Fig. 6B-a, Supplementary Table 1**). Second, we found that the TM mutant of Lyn-GFP (TM-Lyn-GFP; **Fig. 1A**) did not exhibit any detectable longer-lifetime component in the colocalization duration histogram (**Fig. 6B-b, Supplementary Table 1**; *P* = 0.46 against *h*(incidental-by-shift)). These results suggest that (1) the protein moiety of Lyn by itself cannot induce the recruitment, (2) the raft-lipid interaction by itself can induce Lyn recruitment at CD59 clusters, and that (3) when both the Lyn protein moiety and raftophilic myristoyl+palmitoyl chains exist, the lifetime at the CD59 cluster raft appears to be prolonged (could be proven in the future when single-molecule imaging is further improved).

To further examine whether the raft-lipid interaction alone can recruit cytoplasmic saturated-chain-anchored proteins at CD59 clusters, we examined the recruitment of two more artificial molecules with large deletions in their protein moieties, but with preserved lipid binding sites: palpal-N16 GAP43-GFP (raftophilic) and GFP-C5 Rho-gerger (non-raftophilic) (**Fig. 1A**). Palpal-N16 GAP43-GFP exhibited a clear two-component histogram (significant difference from *h*(incidental-by-shift), *P =* 0.0023), with a *τ*_2_ (71 ms) quite comparable to the *τ*_2_’s for Lyn-FG and myrpal-N20(Lyn)-GFP with CD59 clusters (**Fig. 6B-c, Supplementary Table 1**). Meanwhile, GFP-C5 Rho-gerger did not exhibit any detectable *τ*_2_ component (**Fig. 6B-d, Supplementary Table 1**; *P* = 0.97 against *h*(incidental-by-shift)). Taken together, the results obtained with these four designed molecules (**Fig. 6B**) suggest that a raft-lipid interaction without a specific protein-protein interaction could induce the recruitment of cytoplasmic proteins with two saturated chains at CD59 clusters. However, if the protein-protein interaction does exist (Lyn-FG; *τ*_2_ = 80 ms), then it could slightly prolong the colocalization lifetime (myrpal-N20(Lyn)-GFP; *τ*_2_ = 66 ms). In short, the outside-in inter-layer coupling occurs when stabilized CD59-cluster rafts are induced in the outer leaflet, and the out-in transbilayer coupling mechanism is predominantly lipid-based.

### H-Ras is continually and transiently recruited at CD59 clusters in a manner dependent on raft-lipid interactions

Next, we examined the recruitment of fluorescently-labeled H-Ras (FGH-Ras), which is anchored to the PM inner leaflet via two saturated (palmitoyl) chains and an unsaturated (farnesyl) chain covalently conjugated to the C-terminal domain of H-Ras (**Fig. 1A**). Virtually all of the FGH-Ras molecules underwent thermal diffusion, with a diffusion coefficient (in the time scale of 124 ms) of 1.12 ± 0.0017 μm^2^/s (**Fig. 4B**). The FGH-Ras was functional, because it was activated by EGF stimulation (**Fig. S2B**).

The histogram of the colocalization durations of FGH-Ras at CD59 clusters exhibited two clear components (**Fig. 6C-a, Supplementary Table 1**; *P* = 0.029 against *h*(incidental-by-shift); *τ*_2_ = 91 ms), whereas no significant *τ*_2_ component was detected in the histogram for the colocalizations at *non-clustered* CD59 (**Fig. 6C-b, Supplementary Table 1**; *P* = 0.52 against *h*(incidental-by-shift)). After mildly treating the cells with methyl-β-cyclodextrin (MβCD, 4 mM at 37°C for 30 min), the FGH-Ras colocalization with CD59 clusters was strongly suppressed (**Fig. 6C-c**; *P* = 0.41 against *h*(incidental-by-shift)). The strong effect of partial cholesterol depletion supports the critical importance of raft-lipid interactions for the recruitment of lipid-anchored FGH-Ras at CD59 clusters.

Next, we examined the colocalization of GFP-tH (**Fig. 1A**), which lacks the majority of the H-Ras protein moiety (Prior et al., 2001, 2003), with CD59 clusters. The colocalization duration histogram exhibited two clear components (**Fig. 6C-d, Supplementary Table 1**; *P* = 0.027 against *h*(incidental-by-shift); *τ*_2_ = 75 ms vs. 91 ms for the full-length FGH-Ras; non-significant difference). Taken together, these results suggest that the two palmitoyl chains of H-Ras probably mask the effect of the unsaturated farnesyl chain, and thus FGH-Ras’s two palmitoyl chains might work like Lyn-FG’s myristoyl+palmitoyl chains. The *τ*_2_ values are summarized in **Fig. 6D**.

In the present report, we focused on the recruitment of lipid-anchored cytoplasmic signaling molecules, Lyn-FG and FGH-Ras, at CD59-cluster rafts. Since Lyn-FG and FGH-Ras are continually recruited to CD59 clusters, they are considered to be more concentrated within the nano-scale region (on the order of 10 nm) of the CD59-cluster raft. This will enhance the homo- and hetero-interactions of Lyn, H-Ras, and other recruited raftophilic signaling molecules at CD59-cluster rafts. Indeed, we found that the homo-oligomerization of FGH-Ras by crosslinking its FKBP domain by the AP20187 addition could activate FGH-Ras (**Fig. S2**). This result further suggests that the recruitment of Lyn-FG and FGH-Ras at the small cross-sectional area of the CD59-cluster raft, leading to their higher concentrations at CD59 clusters, would have important signaling consequences.

### GM1 clusters formed by antibody-crosslinked CTXB in the PM outer leaflet activate ERK1/2 kinases

To further investigate the *raft-lipid* interactions across the bilayer, we induced clusters of GM1, a prototypical raft-associated glycosphingolipid (ganglioside), in the PM outer leaflet, and examined whether Lyn-FG and FGH-Ras located in/on the inner leaflet could be recruited at GM1 clusters in the outer leaflet. GM1 clusters were induced by applying cholera toxin B subunit (CTXB) conjugated with A633 (dye/protein mole ratio = 0.8), which could bind five GM1 molecules (CTXB-5-GM1; Merritt et al., 1994), and greater GM1 clusters containing an average of 3 CTXB and 15 GM1 molecules (virtually 30 saturated acyl chains) were induced by the further addition of a goat polyclonal anti-CTXB antibody-IgG (Ab-CTXB-GM1 clusters; **Fig. 7A**; see the caption to **Fig. 7B** and **Materials and methods**). We anticipated that all five of the of the GM1 binding sites in CTXB are filled with GM1, since GM1 exists abundantly in the PM outer leaflet of HeLa cells and the two-dimensional collision rate is much higher, as compared with that in three-dimensional space (Grasberger et al., 1986).

**Figure 7.**
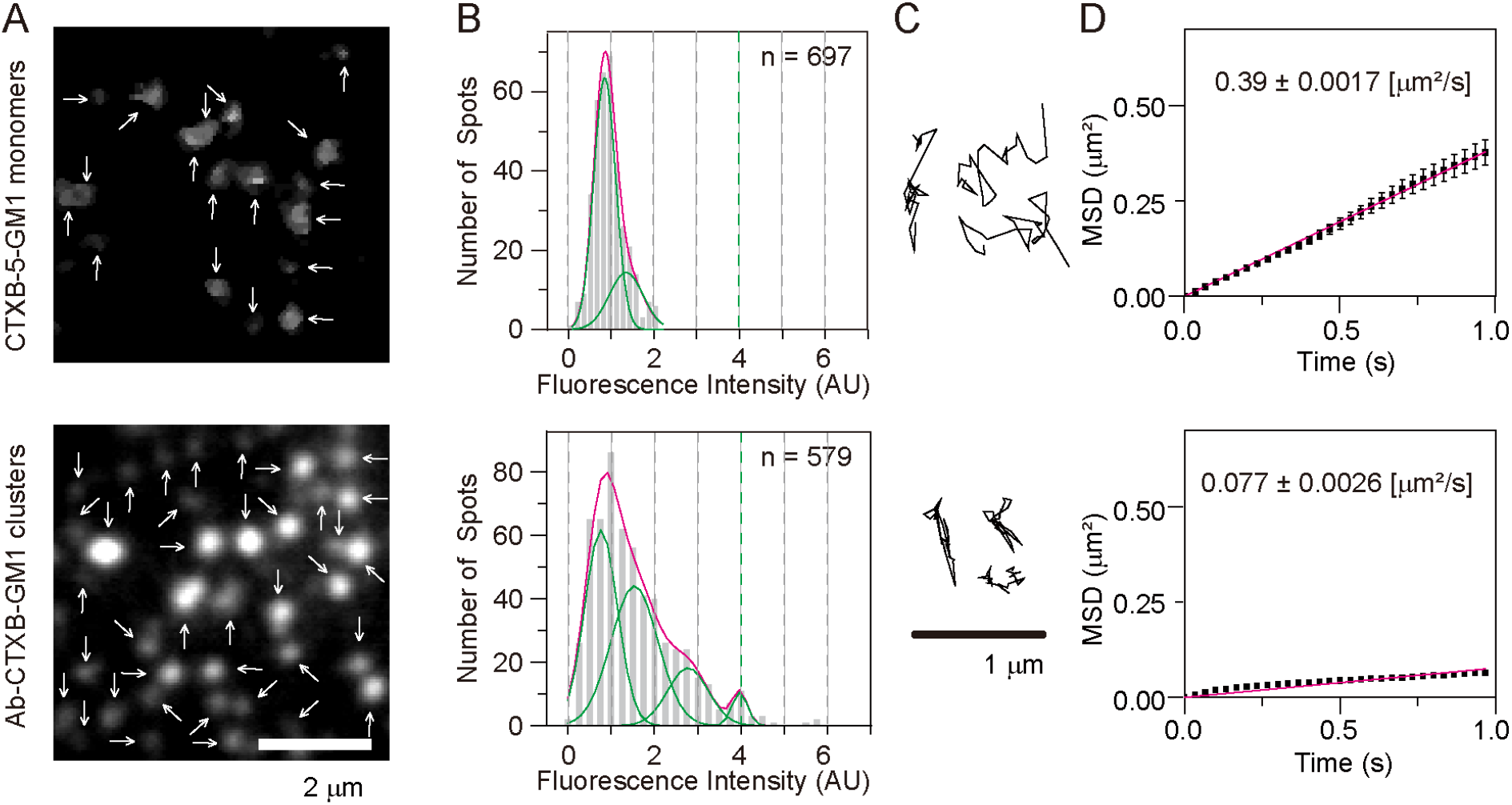
Ab-CTXB-GM1 clusters generated in the PM outer leaflet contained an average of ~15 GM1 molecules and diffused 2.6x slower than CD59 clusters. **(A)** Fluorescence images of non-crosslinked fluorescently-labeled CTXB (which could bind up to five GM1 molecules; A633 conjugated with a dye-to-protein ratio of 0.80; called CTXB-5-GM1; top) and CTXB clusters induced by the further addition of anti-CTXB antibodies (Ab-CTXG-GM1 cluster; bottom), obtained at single-molecule sensitivities. Arrows indicate all of the detected fluorescent spots in each image. **(B)** Histograms showing the distributions of the signal intensities of individual fluorescent spots of A633-labeled-CTXB-5-GM1 (top) and Ab-CTXB-GM1 clusters (bottom). Based on these histograms, we concluded that each Ab-CTXB-GM1 cluster contained an average of ~15 GM1 molecules (see **Materials and methods**), although the distribution would be quite broad. **(C)** Typical trajectories of CTXB-5-GM1 (top) and Ab-CTXB-GM1 clusters (bottom) for 0.2 s, obtained at a time resolution of 6.45 ms. **(D)** Ensemble-averaged MSDs plotted against time, suggesting that in the time scale of 1 s, both CTXB-5-GM1 (top) and Ab-CTXB-GM1 clusters (bottom) undergo effective simple-Brownian diffusion, and the diffusion is slowed by a factor of about five after antibody-induced clustering.

Ab-CTXB-GM1 clusters (30 saturated alkyl chains) diffused with a mean diffusion coefficient of 0.077 μm^2^/s, 5.1x slower than non-crosslinked CTXB-5-GM1 (5 saturated alkyl chains; 0.39 μm^2^/s) (**Fig. 7D**), whereas they diffused 2.6x slower than CD59 clusters (0.20 μm^2^/s; 20 saturated alkyl chains) (**Fig. 2D**). Namely, the average cross-section area of the hydrophobic region of the Ab-CTXB-GM1 cluster would be somewhat greater than that of the CD59 cluster.

The GM1 clusters slowly became entrapped in caveolae; ~9% of the fluorescent spots were colocalized with caveolae at 10 min after the addition of the anti-CTXB antibodies at 27°C (**Fig. S1B**). Therefore, in the present investigation, all of the microscopic observations involving GM1 clusters were made within 10 min after the addition of the antibodies, when most of Ab-CTXB-GM1 clusters were located outside caveolae.

CTXB binding to the cell surface did not trigger the ERK signaling cascade, but when Ab-CTXB-GM1 clusters were induced, the ERK signaling cascade was activated (**Fig. 3**), consistent with the previous observations (Janes et al., 1999; Kiyokawa et al., 2005). We suspect that this is not simply due to the differences in the sizes of the entire CTXB-5-GM1 and Ab-CTXB-GM1 clusters, but rather to those in the local densities of the saturated chains in the CTXB-5-GM1 and Ab-CTXB-GM1 clusters, based on the following reason.

A crystallographic study showed that the five GM1 binding sites in CTXB are all located ~3.7 nm away from adjacent binding sites (**Fig. 1C**; Merritt et al., 1994), and thus the five GM1 molecules (10 saturated chains) are located on a circle with a diameter of ~6.3 nm. Considering the size of the acyl chains (occupying a cross-section of <0.35 nm^2^; i.e., a diameter of <0.33 nm), two adjacent GM1 molecules bound to CTXB will be located >3 nm away from each other, in a space that could accommodate >9 acyl chains, and in the middle of the GM1 binding sites, there would be a circular space with a cross-section of >5 nm in diameter, which could accommodate >25 acyl chains (in the outer leaflet; **Fig. 1C**). Namely, the space between two GM1 molecules with saturated acyl chains bound to a CTXB molecule is much larger than the cross-section of a few lipid molecules; i.e., CTXB induces only sparse GM1 clusters. The observation that CTXB molecules simply bound to the PM cannot trigger the downstream signals is consistent with this consideration: the five GM1 molecules bound to a single CTXB molecule would not provide the threshold densities of saturated lipids necessary to create stable rafts by assembling and keeping cholesterol and saturated chains in CTXB-5-GM1 and excluding unsaturated chains. The five GM1 molecules bound to a single CTXB molecule would not serve as a nucleus to induce raft domains beneath the CTXB molecule (in the outer leaflet of the PM), and hence would fail to recruit signaling molecules that trigger the ERK signaling cascade. This possibility was directly examined in the present study (see the next section; in the case of CD59 clusters, we suspect that due to the long flexible glycochain of GPI, CD59 has reorientation freedom and thus the saturated chains of CD59 in the cluster and the cholesterol, sphingomyelin, and gangliosides recruited from the bulk PM can form a tighter complex beneath the cluster of CD59 protein moieties).

Meanwhile, when the CTXB molecules were crosslinked by anti-CTXB antibodies, since the GM1 molecules are bound near the outer edges of CTXB (Merritt et al., 1994), they would be located very close to the GM1 molecules bound to other CTXB molecules in the Ab-CTXB-GM1 cluster (**Fig. 1C**). These closely associated GM1 molecules could form the stable raft nucleus for recruiting cholesterol and lipids with saturated alkyl chains, recruiting raftophilic signaling molecules, and thus triggering the ERK signaling pathways. Therefore, next we investigated whether CTXB-5-GM1 or Ab-CTXB-GM1 clusters could recruit Lyn-FG and FGH-Ras.

### Lyn and H-Ras are continually and transiently recruited at Ab-CTXB-GM1 clusters in a manner dependent on raft-lipid interactions, but not at CTXB-5-GM1

We directly examined whether single molecules of Lyn-FG and FGH-Ras were recruited at CTXB-5-GM1 and Ab-CTXB-GM1 clusters located in/on the PM outer leaflet. As described in the previous section, CTXB-5-GM1 failed to trigger ERK activation, in contrast to Ab-CTXB-GM1 clusters.

The histogram of the colocalization durations of FGH-Ras at Ab-CTXB-GM1 exhibited two clear components, indicating that Lyn-FG was recruited at Ab-CTXB-GM1 (**Fig. 8A-a, Supplementary Table 2**; *P* = 0.013 against *h*(incidental-by-shift); *τ*_2_ = 110 ms). Meanwhile, no significant *τ*_2_ component was detectable for the colocalizations at CTXB-5-GM1 (**Fig. 8A-b, Supplementary Table 2**; *P* = 0.24 against *h*(incidental-by-shift)). Similarly, FGH-Ras was recruited at Ab-CTXB-GM1 (**Fig. 8B-a, Supplementary Table 2**; *P* = 0.025 against *h*(incidental-by-shift); *τ*_2_ = 97 ms), but not at CTXB-5-GM1 (**Fig. 8B-b, Supplementary Table 2**; *P* = 0.52 against *h*(incidental-by-shift)).

**Figure 8.**
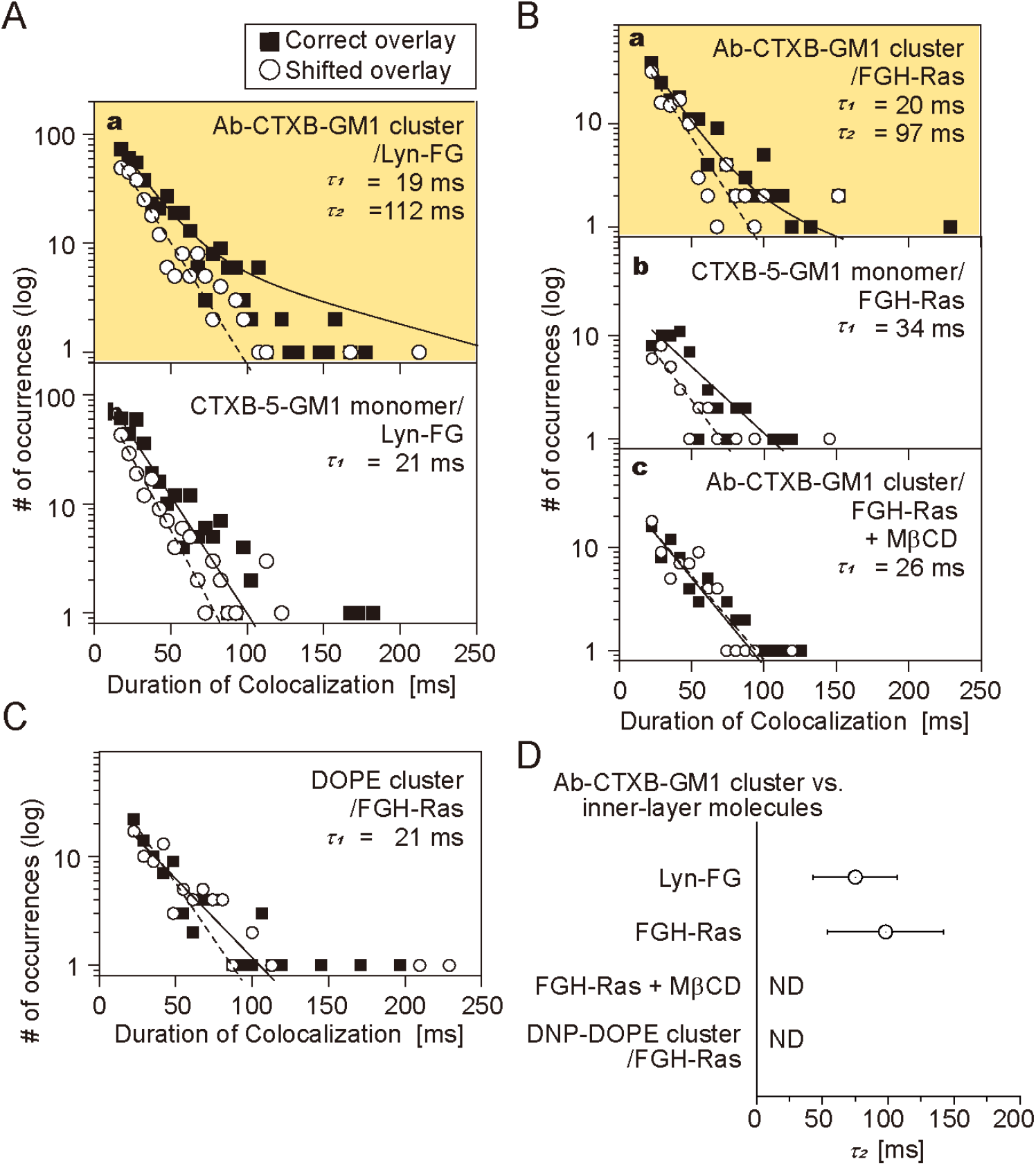
Lyn-FG and FGH-Ras were recruited at Ab-CTXB-GM1 clusters, but not at CTXB-5-GM1. The distributions (histograms) for the colocalization durations are shown. See the caption to **Fig. 6** for details and keys. See **Supplementary Table 2** for statistical parameters.

**(A)** Lyn-FG was recruited at Ab-CTXB-GM1 clusters, but not at CTXB-5-GM1.

**(B)** FGH-Ras was recruited at Ab-CTXB-GM1 clusters, but not at CTXB-5-GM1, and its recruitment at Ab-CTXB-GM1 clusters depended on the PM cholesterol.

**(C)** FGH-Ras was not recruited to DNP-DOPE clusters.

**(D)** Summary of the bound lifetimes (*τ*_2_’s) of Lyn-FG and FGH-Ras at Ab-CTXB-GM1 clusters. ND, the *τ*_2_ component not detected.

Partial cholesterol depletion eliminated the *τ*_2_ component for the FGH-Ras colocalization with Ab-CTXB-GM1 (**Fig. 8B-c, Supplementary Table 2**; *P* = 0.96 against *h*(incidental-by-shift)). Furthermore, when DNP-DOPE, a non-raft reference unsaturated phospholipid, was clustered in the outer leaflet by the addition of anti-DNP antibodies and secondary antibodies (the antibody concentrations were adjusted so that more than 90% of DNP-DOPE clusters became immobile; i.e., the cross section of the DNP-DOPE cluster would be substantially greater than that of Ab-CTXB-GM1), no significant *τ*_2_ component was detected for the FGH-Ras colocalization with DNP-DOPE clusters (**Fig. 8C, Supplementary Table 2**; *P* = 0.39 against *h*(incidental-by-shift)). These results further support the proposal that raft-lipid interactions are essential for the recruitment of cytoplasmic lipid-anchored signaling molecules at Ab-CTXB-GM1, and therefore that the GM1 molecules closely apposed to each other inside the Ab-CTXB-GM1 cluster induce stable raft nuclei by recruiting cholesterol and other raftophilic molecules. The results for *τ*_2_ are summarized in **Fig. 8D**.

### Small clusters of inner-leaflet signaling molecules did not recruit CD59 or GM1 in the outer leaflet

The homo-oligomerization of Lyn-FG and FGH-Ras in the cytoplasm was induced by the addition of AP20187 (dimerizer system developed by Schreiber and then ARIAD; (Schreiber, 1991; Clackson et al., 1998); the presence of a single FKBP molecule in a protein could only create dimers but not oligomers greater than dimers upon AP20187 addition, but the presence of two FKBP molecules in a single protein could induce oligomers (**Fig. 1B bottom**). The average number of Lyn-FG or FGH-Ras molecules in a single cluster was estimated to be ~3 (**Fig. S2C, D** and **Materials and methods**).

The oligomerized FGH-Ras triggered the downstream signaling, as shown by the pull-down assay using the Ras-binding domain of the downstream kinase Raf-1 (**Fig. S2B**), consistent with previous observations (Inouye et al., 2000; Nan et al., 2015). Meanwhile, the oligomerization-induced self-phosphorylation of Lyn-FG was not detected (**Fig. S2A**).

We examined whether CD59 and CTXB-5GM1 located in the PM outer leaflet could be recruited at FGH-Ras or Lyn-FG oligomers induced in the PM inner leaflet, by the addition of AP20187 (**Fig. 9** and **Supplementary Table 3**). No significant recruitment was detectable, indicating that the oligomers of the inner-leaflet lipid-anchored signaling molecules cannot recruit the outer-leaflet raft-associated molecules. This result suggests that although FGH-Ras and Lyn-FG could be transiently recruited to stabilized raft domains, they would only be passengers and not the main molecules for inducing raft domains, probably due to their shorter saturated chains (palmitoyl) and the presence of unsaturated chains. Furthermore, FGH-Ras and Lyn-FG could only be recruited to the outer edges of the raft domains or perhaps the interfaces of the raft and bulk domains. Meanwhile, the lack of CD59 and GM1 recruitment might be due to the smaller sizes (an average of ~3 molecules) of the FGH-Ras and Lyn-FG oligomers.

**Figure 9.**
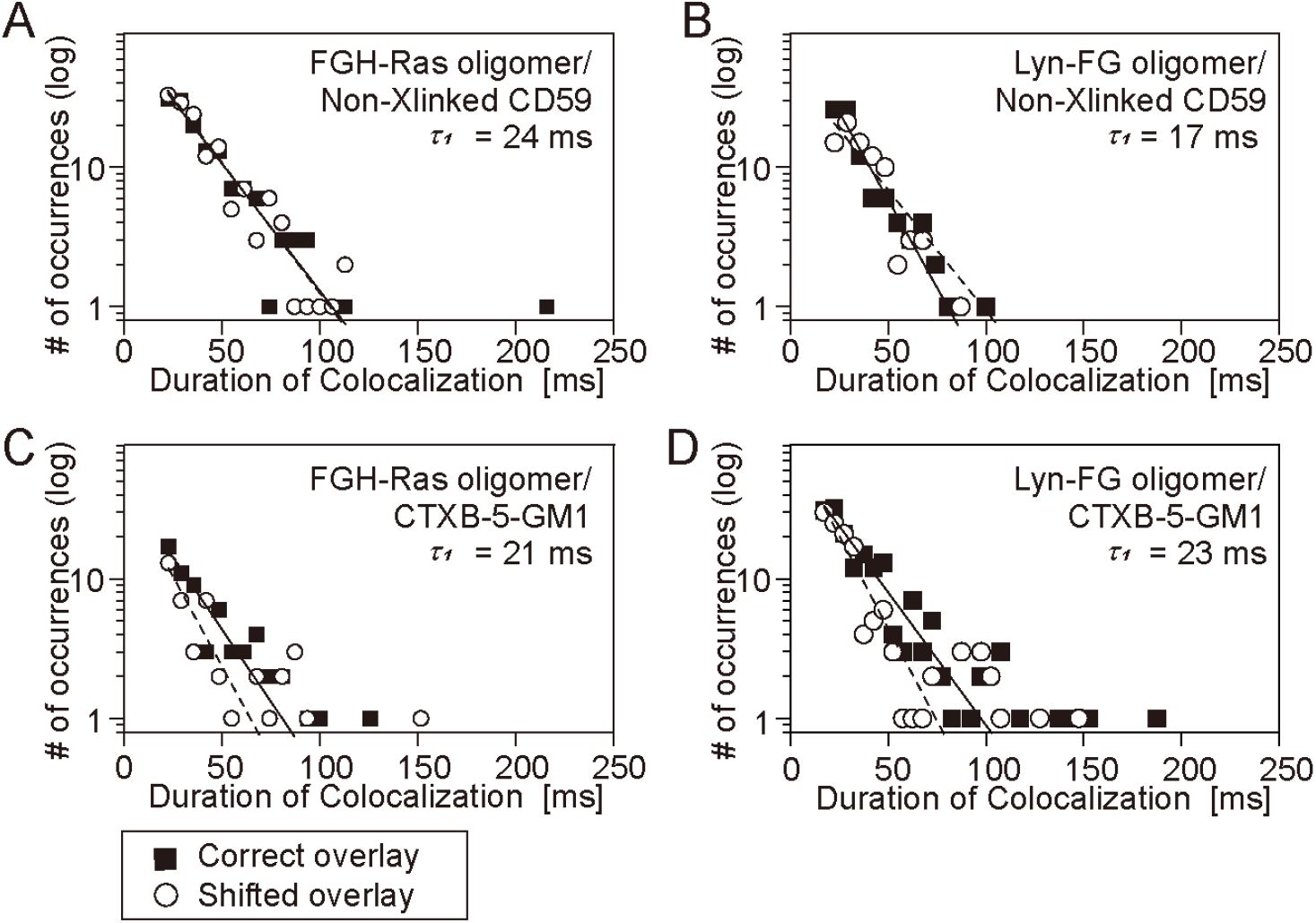
FGH-Ras oligomers and Lyn-FG oligomers induced by the AP20187 addition failed to recruit non-crosslinked CD59 and CTXB-5-GM1. Shown here are the histograms for the durations in which non-crosslinked CD59 and CTXB-5-GM1 located in/on the PM outer leaflet are colocalized with FGH-Ras oligomers and Lyn-FG oligomers artificially induced in the PM inner leaflet by the addition of AP20187. See the caption to **Fig. 6** for details and keys. See **Supplementary Table 3** for statistical parameters. **(A)** Recruitment of non-crosslinked CD59 located in the outer leaflet at the induced FGH-Ras oligomers located in the inner leaflet. **(B)** Recruitment of non-crosslinked CD59 located in the outer leaflet at the induced Lyn-FG oligomers located in the inner leaflet. **(C)** Recruitment of CTXB-5-GM1 located in the outer leaflet at the induced FGH-Ras oligomers located in the inner leaflet. **(D)** Recruitment of CTXB-5-GM1 located in the outer leaflet at the induced Lyn-FG oligomers located in the inner leaflet.

## DISCUSSION

The recruitment of cytoplasmic signaling molecules to small regions in the PM after stimulation is considered to be important for inducing the downstream signaling, since higher concentrations of signaling molecules in small regions would enhance homo- and hetero-interactions, and possibly the formation of transient dimers and oligomers. We indeed found that the homo-oligomerization of FGH-Ras induced by AP20187 activates the downstream signaling of FGH-Ras and H-Ras (**Fig. S2B**). Therefore, in the present research, we extensively studied the recruitment of Lyn, H-Ras, and other lipid-anchored cytoplasmic molecules at CD59-cluster rafts and Ab-CTXB-GM1 clusters.

Our results clearly showed that Ab-CTXB-GM1 clusters of a raftophilic lipid (GM1) formed in the PM outer leaflet can recruit the cytoplasmic lipid-anchored signaling molecules Lyn and H-Ras to the inner leaflet region apposed to the outer-leaflet Ab-CTXB-GM1 clusters. This recruitment was not induced after the PM cholesterol was mildly depleted or when the unsaturated lipid (DNP)-DOPE was clustered in the outer leaflet. These results unequivocally demonstrate that the cytoplasmic lipid-anchored signaling molecules Lyn and H-Ras can be assembled at the stabilized raft-lipid clusters formed in the outer leaflet by raft-lipid interactions. The involvement of TM proteins in the recruitment process would be quite limited, since (1) even cytoplasmic lipid-anchored molecules after the deletions of the majorities of their protein moieties were recruited at Ab-CTXB-GM1 clusters, (2) their dwell lifetimes at CD59 clusters and Ab-CTXB-GM1 clusters were very similar to those of Lyn-FG and FGH-Ras, and (3) the recruitment of FGH-Ras depended on the PM cholesterol level. Of course, this does not rule out the specific interactions of GPI-ARs with TM proteins, as co-receptors (Klein et al., 1997; Wang et al., 2002; Zhou, 2019).

To summarize the sequence of events in CD59 signaling (**Fig. 10**), first, the stable CD59-cluster rafts in the outer leaflet are induced by the clustering of raftophilic CD59 molecules by extracellular stimulation, such as MACC binding. The stabilized raft domains tend to last for durations on the order of 10s of minutes (Suzuki et al., 2007b; Suzuki et al., 2012), whereas their constituent molecules, such as the gangliosides, tend to stay there only for 50 ms and turnover quickly, continually exchanging with those located in the bulk PM region (Komura et al., 2016). Second, at the signal-induced stabilized CD59-cluster raft domains, the raftophilic cytoplasmic signaling molecules, Lyn and H-Ras, are recruited by raft-lipid interactions with lifetimes on the order of 0.1 s (**Figs. 6 and 8**); i.e., each molecule stays at the CD59-cluster raft quite transiently. However, since many molecules would continually arrive one after another, and since each raft domain can accommodate several hundred lipid molecules (when the raft radius is 10 nm, each leaflet within the raft can accommodate ~500 phospholipids), many cytoplasmic raftophilic signaling molecules could be dynamically concentrated in the small cross-sectional area beneath the CD59-cluster raft, leading to locally enhanced molecular interactions.

**Figure 10.**
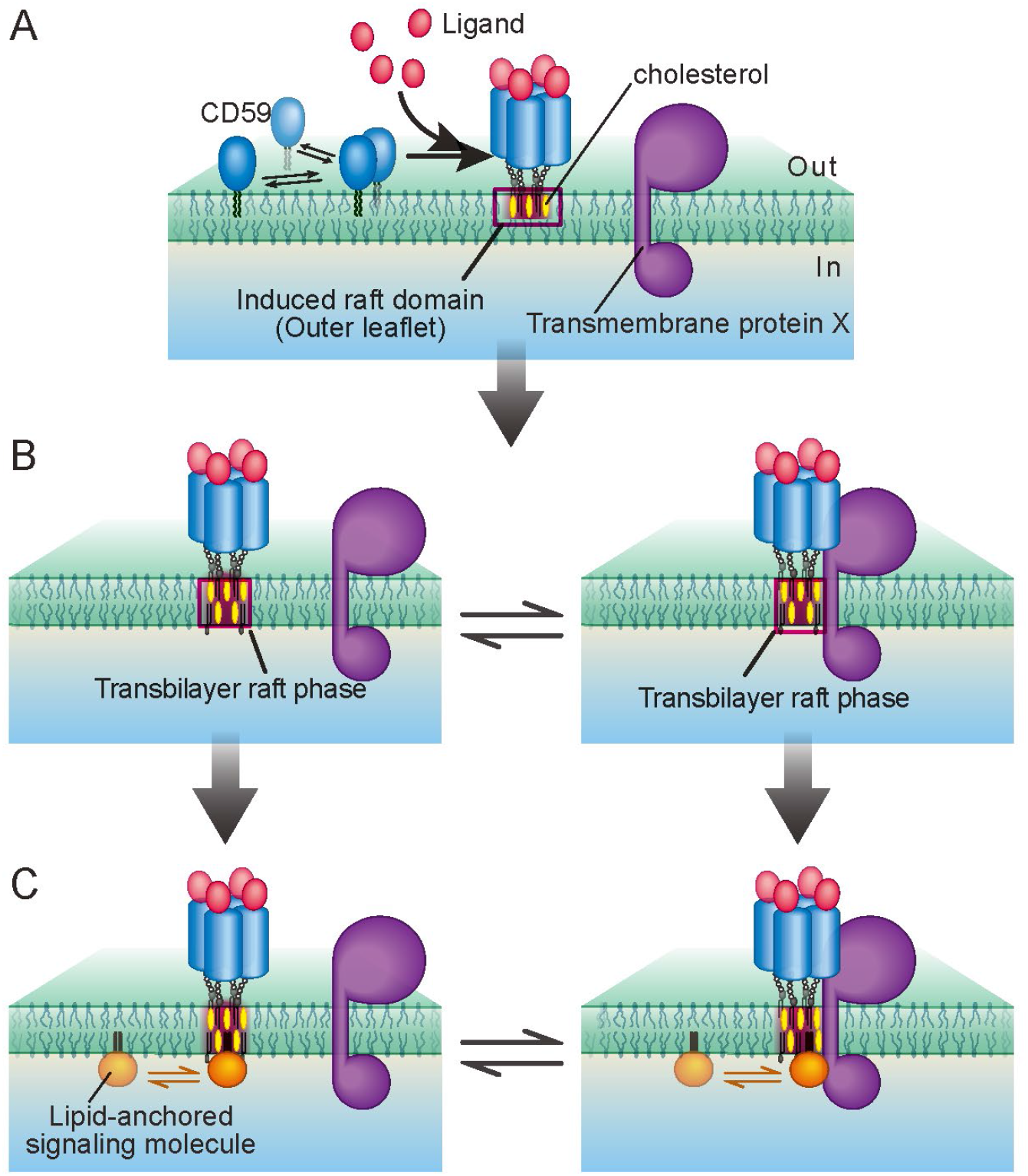
Schematic model showing the CD59 signal transduction mediated by the transbilayer raft phase, which recruits lipid-anchored signaling molecules at the ligated, stabilized CD59-cluster domains in the PM outer leaflet, inducing enhanced interactions of recruited molecules. **(A)** First, the ligand binding triggers the conformational changes of CD59, which in turn induce CD59 clustering, creating stable DC59-cluster signaling rafts. If GM1 is clustered closely, then stable GM1-cluster rafts will be produced. **(B)** Then, the transbilayer raft phase is induced by the CD59-cluster raft, by involving molecules in the inner leaflet, recruiting cholesterol and molecules with saturated alkyl chains (left) and also excluding molecules with unsaturated alkyl chains. An as-yet-unknown TM protein(s) X, which has affinities to raft domains, might also be recruited to the transbilayer raft phase (right; The recruitment of X could be enhanced by specific protein-protein interactions with the ligated CD59 exoplasmic protein domain). **(C)** Finally, cytoplasmic lipid-anchored signaling molecules, such as H-Ras and Lyn, are recruited to the transbilayer raft phase in the inner leaflet by the raft-lipid interaction (left). This could be enhanced by the protein-protein interaction with the transmembrane protein X (right). Although the residency times of the inner-leaflet signaling molecules beneath the CD59 cluster may be limited, since many molecules will be recruited there one molecule after another, interactions of two or more species of cytoplasmic signaling molecules will occur efficiently beneath the CD59-cluster raft. This way, the transbilayer raft phase induced by the stabilized CD59-cluster raft would function as an important signaling platform.

Let us assume that the sizes of the stabilized CD59-cluster rafts and Ab-CTXB-GM1 cluster rafts are in the ranges of 20 – 100 nm in diameter (**Figs. 1B, C, 2 and 7**) and the diffusion coefficient of the lipid-anchored signaling molecules is ~1 μm^2^/s (**Fig. 4**). Then, these signaling molecules would stay in the 20 – 100 nm region in the bulk PM for only 0.03 – 0.63 ms. However, they remained in the stabilized raft domains for 80 – 110 ms (**Figs. 6 and 8**); i.e., the dwell lifetimes were prolonged by a factor of 200~2,000, which is a large factor. Namely, the dwell lifetimes in the range of 80 – 110 ms might appear to be short, but in fact, Lyn-FG, FGH-Ras, and other raftophilic lipid-anchored molecules exhibited extremely prolonged dwell lifetimes beneath the stabilized raft domains in the outer leaflet. Such extreme prolongation would not be possible by simple interactions of the lipids in the inner leaflet with the lipids in stabilized raft domains in the outer leaflet.

Since the micron-scale transbilayer raft phases have been detected in artificial bilayer membranes (Collins and Keller, 2008; Blosser et al., 2015), we propose that nano-scale transbilayer raft phases are induced in both leaflets by the stabilized raft domains initially formed in the outer leaflet, and that when cytoplasmic raftophilic lipid-anchored signaling molecules arrive at the transbilayer raft phases, they tend to be trapped in the inner-leaflet part of the transbilayer raft phase, exhibiting dwell lifetimes on the 0.1-s order. Namely, *transbilayer raft-lipid interactions* would only make sense when *cooperative* lipid interactions occur due to the formation of the transbilayer raft phase. Indeed, raft domains are generally considered to form by the cooperative interactions of saturated alkyl chains and cholesterol, as well as by their cooperative exclusions from the bulk PM enriched in unsaturated alkyl chains. Therefore, we propose that the nano-scale transbilayer raft phases would be induced at stabilized rafts initially formed in the outer leaflet. We further propose that the transbilayer raft phases induced by the stimulation-triggered GPI-AR-cluster rafts would act as a key signaling platform for the engaged GPI-AR clusters, and that the formation of the transbilayer raft phase would be a general mechanism for GPI-AR signal transduction.

This recruitment mechanism based on the transbilayer raft phase appears to suggest the lack of specificity in the cytoplasmic signaling, without any dependence on the GPI-AR species. However, since different GPI-ARs would form signaling cluster rafts with a variety of sizes, closeness of the saturated acyl chains within the cluster (as found here for GM1 clusters induced by CTXB), and the TM protein species with which different GPI-ARs interact would vary (Suzuki et al., 2012; Zhou, 2019), the GPI-AR-cluster rafts formed in the outer leaflet could induce transbilayer raft phases with distinct properties. These transbilayer raft phases could recruit a variety of lipid-anchored signaling molecules with differing efficiencies, thus triggering various downstream signaling cascades with different strengths; i.e., the relative activation levels among the many intracellular signaling cascades triggered by GPI-ARs would vary depending on the GPI-AR species.

## Materials and methods

### Improved camera systems for simultaneous, dual-color, single-molecule imaging in living cells at enhanced time resolutions of 5 and 6.45 ms

The major improvement of our single-molecule imaging station from the previously published version (Koyama-Honda et al., 2005; Komura et al., 2016; Kinoshita et al., 2017) was the employment of two camera systems that allow higher frame rates. With an increase in the frame rate of the camera system, we employed lasers with higher outputs (see the next paragraph). The two camera systems both employed two-stage microchannel plate intensifiers (C8600-03, Hamamatsu). In one camera system, the image intensifier was lens-coupled to an EM-CCD camera (Cascade 650, Photometrics), which was operated at 155 Hz (6.45 ms/frame), with a frame size of 653 x 75 pixels (38.9 x 4.46 μm^2^ for a total of 240x magnification). In the other camera system, the image intensifier was fiber-coupled, with a 1.6:1 tapering, to a CCD camera (XR/MEGA-10ZR, Stanford Photonics) cooled to −20°C and operated at 200 Hz (5 ms/frame), with a frame size of 640 x 160 pixels (27.1 x 6.75 μm^2^ for a total of 240x magnification).

The bottom PMs of the cells cultured on glass-bottom dishes were observed by a home-built objective lens-type TIRF microscope with two detection arms for simultaneous two-color single-molecule imaging, as described previously (Koyama-Honda et al. 2005). The temperature of the sample and the microscope was maintained at 27 ± 1°C. The cells were illuminated simultaneously by a 488-nm laser (for GFP, SAPPHIRE 488-20, Coherent) and a 594-nm laser (for A633, 05-LYR-173, Melles Griot). Fluorescence signals from GFP and A633 were split into the two detection arms by using a dichroic mirror at 600 nm (600DCXR, Chroma), and further isolated by interference filters (HQ535/70 for GFP and HQ655/100 for A633, Chroma). The fluorescence image in each arm was projected onto the photocathode of the image intensifier in the camera system described above (the same cameras were employed for the two channels).

### Colocalization detection and evaluation of colocalization lifetimes

For the colocalization analysis, GFP-trajectories longer than 19 frames and A633-trajectories longer than 29 frames were used. The colocalization of an A633 spot with a GFP spot was defined as the event where the two fluorescent spots, representing A633 and GFP molecules, became localized within 150 nm from each other. This is a distance at which an exactly colocalized molecule is detected as colocalized at probabilities greater than 90%, using the Cascade 650 camera operated at 155 Hz, and higher probability was achieved using the XR/MEGA-10ZR camera operated at 200 Hz (Koyama-Honda et al., 2005).

A colocalization distance of 150 nm is much greater than the molecular scale, and therefore, in addition to colocalization due to specific molecular binding, events where molecules incidentally encounter each other within a distance of 150 nm, termed ‘incidental colocalizations’, can occur. However, as described in the Results section, non-associated molecules may track together by chance over a short distance, but the probability of moving together for multiple frames is small, and therefore longer colocalizations imply the binding of two molecules.

In the analysis of colocalization durations, those as short as one or two frames were neglected to avoid higher frequency noise. Likewise, if two colocalization events are separated by a gap of one or two frames, then they are linked and counted as a single longer colocalization event. To obtain the histogram of incidental colocalization durations, the image obtained in the longer-wavelength channel (A633) was shifted toward the right by 20 pixels (1.0 and 1.19 μm, depending on the camera) and then overlaid on the image obtained in the GFP channel (“shifted overlay”). The histogram of the incidental colocalization durations was called *h*(incidental-by-shift). We found *h*(incidental-by-shift) could effectively be fitted by a single exponential decay function, using nonlinear least-squares fitting by the Levenberg-Marquardt algorithm provided in the OriginPro software, and the decay time constant was called the incidental colocalization lifetime, *τ*_1_ (**and 8**).

Meanwhile, the distribution of the colocalization durations for correctly overlaid A633 and GFP images (“Correct Overlay”) was obtained, and we found that some of the histograms (such as that for Lyn-FG vs. CD59 clusters) could be fitted with the sum of two exponential functions with a decay time constant *τ*_1_’ and the other longer time constant *τ*_2_ (**Figs. 6 and 8**). The *τ*_1_’ component was considered to represent the duration of incidental colocalization, and thus *τ*_1_’ = *τ*_1_. Therefore, in the following, we describe *τ*_1_’ simply as *τ*_1_.

The *τ*_2_ component of the histogram was considered to describe the colocalization durations, including the durations of true molecular interactions (*τ*_B_). Here, we propose that the binding duration *τ*_B_ can be approximated by *τ*_2_, which can be directly determined from the histogram. As described previously (Kasai et al., 2018), the duration *τ*_2_ would be the sum of (1) the duration between the incidental encounter and actual molecular binding, (2) the duration of molecular binding (*τ*_B_), and (3) the duration between the dissociation of two molecules and separation by >150 nm. Therefore, the mathematical function to describe the histogram for the colocalization durations including the molecular binding would be exp(-*t*/*τ*_B_) convoluted with the histogram *h*(incidental-by-shift), which is proportional to exp(-*t*/*τ*_1_) (*t*; time) at the present experimental accuracies (**Figs. 6 and 8**). Here, we are assuming simple zero-order kinetics for the release of lipid-anchored cytoplasmic molecules from the CD59-cluster rafts (and thus the binding duration distribution is proportional to exp(-*t*/*τ*_B_)). The result of the convolution of an exponential function with another exponential function is well known, and the convoluted function is the sum of these two exponential functions (exp(-*t*/*τ*_1_) and exp(-*t*/*τ*_B_)). Therefore, the entire histogram is the sum of the histogram for simple close encounters, *h*(incidental-by-shift), which has the form of exp(-*t*/*τ*_1_), and the histogram for the colocalization events that include molecular interactions, and is expressed by the sum of exp(-*t*/*τ*_1_) and exp(-*t*/*τ*_B_). Meanwhile, as described, some of the experimentally obtained histograms (such as that for Lyn-FG vs. CD59 clusters) could be fitted with the sum of two exponential functions with the decay time constant *τ*_1_ and the other longer time constant *τ*_2_ (**and 8**). Therefore, we find *τ*_B_ = *τ*_2_. Namely, the longer time constant *τ*_2_ obtained from the fitting represents the binding duration (**Figs. 6D and 8D**).

For the actual two component fitting for the histograms of the correctly overlaid images, the exponential lifetime for the faster decay function was fixed at the *τ*_1_ value determined from the histogram of the shifted overlay *h*(incidental-by-shift), and then the fitting with the sum of two exponential functions was performed. For some intracellular signaling molecules, the second component was undetectable, indicating that the colocalization did not take place. Throughout this report, the Brunner-Munzel test was used for the statistical analysis, and its result as well as the mean, SEM, the number of conducted experiments, and all other statistical parameters are summarized in **Supplementary Tables 1-3**.

However, due to the problem of the signal-to-noise ratios, the actual estimation of *τ*_2_ involved quite large errors. Accordingly, in the present study, we paid more attention to whether the duration histogram could be represented by a single exponential decay function or the sum of two exponential decay functions.

### Plasmid generation

The cDNA encoding two tandem FKBPs (FKBP2) was obtained from the pC4-Fv1E vector (ARGENT Regulated Homodimerization Kit, ARIAD Pharmaceuticals), and subcloned into the pTRE2hyg vector (including a tetracycline-responsible promoter, Takara Bio) with the cDNA encoding GFP-H-Ras (a kind gift from A. Yoshimura; Murakoshi et al., 2004) to produce FKBP2-GFP-H-Ras (FGH-Ras). The cDNA encoding Lyn was obtained from RBL-2H3 cells and subcloned into the pTRE2hyg vector with the cDNA encoding FKBP2 and EGFP (derived from pEGFP-N2; Clontech) to produce Lyn-FKBP2-GFP (Lyn-FG). The cDNA encoding EGFP was subcloned with the signal sequence 5’-ggg tgc ctt gtc ttg tga-3’ for the geranylgeranyl modification (CAAX) into the pTRE2hyg vector to produce GFP-C5 Rho-gerger. The cDNAs encoding myrpal-N20Lyn-GFP, palpal-N16GAP43-GFP, and GFP-tH were constructed as described previously (Pyenta et al., 2001; Zacharias et al., 2002; Prior et al., 2003). The cDNA encoding TM-Lyn-GFP was generated by linking the cDNA sequence for the signal peptide derived from low density lipoprotein receptor (LDLR) to the T7-tag sequence, the transmembrane domain of the LDLR sequence, the cDNA encoding Lyn with a deletion of the N-terminal six amino acids (myr-pal modification site), and then to the GFP sequence, and subcloning the produced cDNA sequence into the pTRE2hyg vector. The Cavelin1-GFP vector and GST-RBD vector were generous gifts from T. Fujimoto (Kogo and Fujimoto, 2000) and A. Yoshimura (Murakoshi et al., 2004), respectively.

### Cell culture, transfection, and expression of chimeric molecules

HeLa Tet-Off cells and Tet-On cells (Clontech) were maintained in Eagle’s minimum essential medium (Invitrogen) supplemented with 10% FBS (Sigma), and transfected with each plasmid using LipofectAMINE Plus (Invitrogen). HeLa Tet-Off cells stably expressing FGH-Ras, and HeLa Tet-On cells stably expressing Lyn-FG, myrpal-N20Lyn-GFP, TM-Lyn-GFP, Palpal-N16GAP43-GFP, GFP-C5 Rho-gerger, and GFP-tH were selected in medium containing 0.2 mg/ml hygromycin, and positive clones were captured with micropipettes. The vector encoding Cavelin1-GFP was transfected using LipofectAMINE Plus, and the protein was transiently expressed in HeLa Tet-On cells. Before single-molecule observations, HeLa cells were replated on 12-mmΦ glass-bottom culture dishes (Iwaki) and cultured for 2~3 days. The medium for the FGH-Ras expressing HeLa Tet-Off cells contained 2 μg/ml doxycycline (Dox, ICN Biomedicals) to reduce the expression of recombinant molecules to levels suitable for single-molecule observations. The medium for Tet-On cells expressing GFP-fusion proteins did not contain Dox because, even without Dox-induced expression, the expression levels were sufficiently high for single-molecule observations. For the western blotting and immunostaining of Lyn-FG, its expression levels were enhanced by incubating the Lyn-FG expressing HeLa Tet-On cells in medium supplemented with 2 μg/ml Dox for 24 h before the subsequent experiments.

### Fluorescence labeling and crosslinking of CD59, GM1, and DNP-DOPE

The anti-CD59 antibody IgG was purified from the supernatant of the culture medium of the mouse hybridoma MEM43/5 cell line (provided by V. Horejsi; Stefanova et al., 1991), and the anti-CD59 Fab was prepared by papain digestion of anti-CD59 IgG, followed by protein G column chromatography. The dye/protein mole ratios of the A633 conjugates with anti-CD59 Fab, anti-CD59 IgG, anti-DNP IgG, and CTXB were 0.3, 0.6, 1.4, and 0.8, respectively.

To fluorescently visualize CD59 without crosslinking, the cells were incubated with 0.14 μg/ml anti-CD59 Fab-A633 in Hanks’ balanced salt solution buffered with 2 mM PIPES at pH 7.4 (P-HBSS), at 27°C for 3 min. To generate CD59 clusters, the cells were first incubated with 0.5 μg/ml anti-CD59 IgG-A633 in P-HBSS at 27°C for 3 min, and then with 1.8 μg/ml anti-mouse-IgG antibodies produced in goat (ICN Biomedical) at 27°C for 10 min. To label GM1, cells were incubated with 1 nM CTXB-A633 in P-HBSS at 27°C for 2 min, which could crosslink up to five GM1 molecules. To generate larger GM1 clusters, after the GM1 labeling with CTXB-A633, the CTXB-A633 was further crosslinked by the addition of goat anti-CTXB antibodies (Calbiochem-Novabiochem), diluted 1/100 with P-HBSS, at 27°C for 10 min.

DNP-DOPE was synthesized essentially as described (Murase et al., 2004). Briefly, after conjugating 2,4-dinitrophenyl-*N*hydroxysuccinimide ester (DNP-NHS, Bayer Schering Pharma) to the amine group of 1,2-dioleoyl-*sn*-glycero-3-phosphoethanolamine (DOPE, Avanti Polar Lipid), DNP-DOPE was purified by silica gel thin-layer chromatography and dissolved in methanol. For observing monomeric DNP-DOPE in the PM, the cells were first incubated with the direct addition of 1 μl of 1 mM DNP-DOPE (in methanol), and then the DNP-DOPE incorporated in the PM was labeled by incubating the cells in HBSS containing 5 nM A633-anti-DNP half-IgG and 1% BSA at 27°C for 3 min. To generate DNP-DOPE clusters in the PM, the cells in P-HBSS were first incubated with 1 μM DNP-DOPE at 27°C for 15 min, followed by the incubation with 100 nM A633-anti-DNP IgG in P-HBSS containing 1% BSA at 27°C for 2 min, then with 170 nM goat anti-rabbit IgG (Cappel) in the same buffer at 27°C for 15 min.

### Estimation of the cluster sizes of CD59, GM1, Lyn-FG, and FGH-Ras

The signal intensities of individual fluorescence spots representing one or more molecules on the PM were estimated by TIRF microscopy, as described previously (Iino et al., 2001). Briefly, the fluorescent signal intensities of 600 nm × 600 nm areas (8-bit images in an area of 12 × 12 pixels), each containing a single spot, were measured. The background intensity estimated in adjacent areas was always subtracted. Histograms were fitted with a multi-peak Gaussian by using Origin5 (OriginLab Corp.). In the case of CD59 clusters (**Fig. 2B**, bottom), the histogram was fitted with the sum of 5 Gaussian functions, using the initial values for the means of *m*, 2*m*, 3*m*, 4*m*, and 5*m* and those for the standard deviations of σ, 2^1/2^ σ, 3^1/2^σ, 2σ, and 5^1/2^σ, respectively, where *m* and σ are the mean signal intensity and standard error for the spots representing single A633-Fab molecules adsorbed on the coverslip, with a certain range limitation for the value of each parameter. This provided a ratio of the five Gaussian integrated components of 18:31:31:18:2.

However, since the dye-to-protein mole ratio (D/P ratio) of anti-CD59 IgG-Alexa was 0.6; i.e., 55% of CD59 molecules are not fluorescently labeled (according to the Poisson distribution), this ratio does not represent the true distribution of the sizes of CD59 clusters (in terms of the number of IgG-Alexa molecules in a cluster: for example, a cluster of 3 CD59 proteins might exist without any fluorescence signal). From the Poisson distribution of a mean D/P ratio of 0.6, the distributions of the molecules with true D/P ratios of 0, 1, 2, and 3 are calculated to be 55:33:9.9:2.0, respectively. We simplified this ratio to 6:3:1:0, and based on this distribution, the signal intensity distribution of real CD59 *N*mers was calculated for the *N* values of 3, 4, 5, and 6. When *N* = 5 (pentamers), the fractions of the fluorescent dye molecules in the fluorescent spots with the mean signal intensities of *m, 2m, 3m, 4m,* and *5m* became 13:40:30:16:1, respectively, which are closest to the observed ratio of 18:31:31:18:2. Therefore, although dimers, trimers, tetramers, hexamers and so on must exist, we believed that CD59 pentamers are the most frequent CD59 clusters. However, in the present study, the fluorescent label was not on CD59, but on the anti-DC59 antibody IgG, and since the efficiency of divalent antibody binding to two CD59 molecules is probably very high due to the two-dimensionality of the CD59 spatial distribution on the PM (Grasberger et al., 1986), we concluded that the most frequently formed CD59 clusters consisted of 10 CD59 molecules.

In the case of Ab-CTXB-GM1 clusters (**Fig. 7B**), the histogram could be fitted with a sum of 4 Gaussian functions, providing a ratio of 40:42:15:3 for the four integrated components. The D/P ratio for A633-CTXB was 0.8. From the Poisson distribution of a mean D/P ratio of 0.8, the distribution of the molecules with true D/P ratios of 0, 1, 2, and 3 is calculated to be 45:36:14:4, respectively. We simplified this ratio to 5:4:1:0, and based on this distribution, the signal intensity distribution of the *N*mers of CTXB (Ab-CTXB-GM1 clusters) was calculated for the *N* values of 1, 2, 3, and 4. When *N* = 3 (trimers of CTXB), the fractions of the fluorescent dye molecules in the fluorescent spots with the mean signal intensities of *m, 2m, 3m,* and *4m* became 37:37:23:3, respectively, which are closest to the observed ratio of 40:42:15:3. As in the case with the CD59 clusters, although dimers, tetramers, pentamers, and hexamers and so on must exist, we believe that the most frequent Ab-CTXB-GM1 clusters are those based on CTXB trimers. Using the same argument as for CD59 clusters, each CTXB is expected to be bound by 5 GM1 molecules, and thus we expect that each Ab-CTXB-GM1 cluster usually contains 15 GM1 molecules. This number is quite comparable to the presence of 10 CD59 molecules in a CD59-cluster raft.

### Crosslinking FGH-Ras and Lyn-FG on the cytoplasmic surface of the PM

AP20187, containing two binding sites for the FKBP protein, and AP21998, containing a single binding site for the FKBP protein; i.e., a control molecule for AP20187, were obtained from ARIAD and stored in ethanol, according to the manufacturer’s recommendations. The FKBP fusion proteins FGH-Ras and Lyn-FG, which contain two tandem FKBP moieties in a single molecule, were crosslinked by incubating the cells with 10 nM AP20187 in the culture medium at 27°C for 10 min.

### Evaluating the activation (biological function) of FGH-Ras by crosslinking with AP20187 or the addition of EGF

GST-RBD was expressed in *E. coli* and purified with glutathione-Sepharose beads (Amersham). HeLa Tet-Off cells expressing FGH-Ras (30% confluence in a 10 cm dish) were cultured without serum 24~48 h before the assay. After treating the cells with 10 nM AP20187 or AP21998 for 10 min or 20 nM EGF for 5 min at 37°C, the cells were extracted for 10 min with 1 ml ice-cold buffer, containing 120 mM NaCl, 10% glycerol, 0.5% Triton X-100, 10 mM MgCl2, 2 mM EDTA, 1 μg/ml aprotinin, and 1 μg/ml leupeptin, buffered with 20 mM N-(2-hydroxyethyl)piperazine-N’-ethanesulfonic acid at pH 7.5 (assay buffer). After brief centrifugation (15,000 rpm for 10 min), 20 μl of an RBD-GST/glutathione-Sepharose bead suspension, prepared as described previously (de Rooij and Bos, 1997; Sydor et al., 1998), was added to the supernatant, and the mixture was incubated at 4°C for 1 h. The activated H-Ras molecules would become bound to the RBD-GST-conjugated beads in this process. The beads were then precipitated by centrifugation at 15,000 rpm for 1 min, washed three times with assay buffer, and then after the final centrifugation, the pellet was mixed with 50 μl SDS sample buffer. After SDS-PAGE, western blotting was conducted with mouse anti-Ras antibodies (BD Transduction Laboratories).

### Evaluating the activation (biological function) of Lyn-FG by crosslinking with AP20187 or by antigen stimulation using rat basophilic leukemia (RBL) cells

RBL-2H3 cells expressing Lyn-FG (30% confluence in a 10 cm dish) were cultured without serum 24~48 h before the assay. The high-affinity Fc_ε_ receptor (Fc_ε_RI) was bound by anti-DNP IgE by incubating the cells with 1 μg/ml anti-DNP IgE (SIGMA) overnight. The cells were then incubated with 10 nM AP20187 or AP21998 for 10 min or 100 ng/ml DNP-BSA (Sigma) at 37°C for 60 min, and then extracted on ice for 10 min with 0.3 ml ice-cold extraction buffer, containing 1% NP-40, 0.25 % sodium deoxycholate, 150 mM NaCl, 1 mM EDTA, 0.1 % protease inhibitor mix (Sigma), and phosphatase inhibitors (1 mM Na2VO3 and 1 mM NaF), buffered with 50 mM Tris-HCl at pH 7.4. After brief centrifugation (15,000 rpm for 10 min), the supernatant was mixed with 0.1 ml 4x sample buffer and incubated at 95°C for 5 min. The proteins in the extract were separated by SDS-PAGE, and then western blotting was performed with rabbit anti-pY418 antibodies (BioSource International) and rabbit anti-Lyn antibodies (Santa Cruz).

## Online supplemental material

Fig. S1 shows that less than 10% of CD59-cluster rafts and Ab-CTXB-GM1 clusters became trapped in caveolae within 10 min after their induction. Fig. S2 shows that small clusters of FGH-Ras and Lyn-FG formed in the inner-leaflet triggered signal transduction.

Supplementary Tables 1, 2, and 3 summarize the colocalization lifetimes (*τ*_1_, *τ*_2_) and statistical parameters for the molecular recruitment. Video 1 shows typical single-molecule image sequences of a transient colocalization event of a CD59 cluster and a single Lyn-FG molecule, recorded at a 6.45-ms resolution (155 Hz).

## Acknowledgements

We thank Prof. A. Yoshimura of Keio University School of Medicine for the kind gifts of the cDNAs encoding GFP-H-Ras and GST-RBD, Prof. T. Fujimoto of Nagoya University School of Medicine for the cDNA encoding cavelin1-GFP, and Prof. V. Horejsi of the Academy of Sciences of the Czech Republic for the mouse hybridoma MEM43/5 cell line. This work was supported in part by Grants-in-Aid for scientific research from the Japan Society for the Promotion of Science (Kiban C to I. Koyama-Honda [JP17K07302], Kiban B to T.K. Fujiwara [16H04775], Kiban C to R.S. Kasai [17K07333], Kiban B to K.G.N. Suzuki [18H02401], and Kiban S to A. Kusumi [16H06386]) and the program of Exploratory Research for Advanced Technology Organization (ERATO) of the Japan Science and Technology Agency (JST) (to Prof. Noboru Mizushima of the University of Tokyo [JPMJER1702]). WPI-iCeMS of Kyoto University is supported by the World Premiere Research Center Initiative (WPI) of MEXT.

The authors declare no competing financial interests.

Author contributions. I. Koyama-Honda performed a large majority of the single fluorescent-molecule tracking experiments. I. Koyama-Honda, E. Kajikawa, and H. Tsuboi conducted biochemical and cell biological experiments. I. Koyama-Honda, T.K. Fujiwara, R.S. Kasai, and A. Kusumi developed and built the single-molecule imaging station that can operate at higher frequencies. T.K. Fujiwara, R.S. Kasai, and T.A. Tsunoyama developed the analysis software. I. Koyama-Honda, K.G.N. Suzuki, and A. Kusumi conceived and formulated the project. I. Koyama-Honda, T.K. Fujiwara, R.S. Kasai, K.G.N. Suzuki, and A. Kusumi evaluated the obtained data and extensively discussed experimental plans during the entire course of this research. I. Koyama-Honda and A. Kusumi wrote the manuscript, and all authors participated in revisions.

---

**Figure S1.**
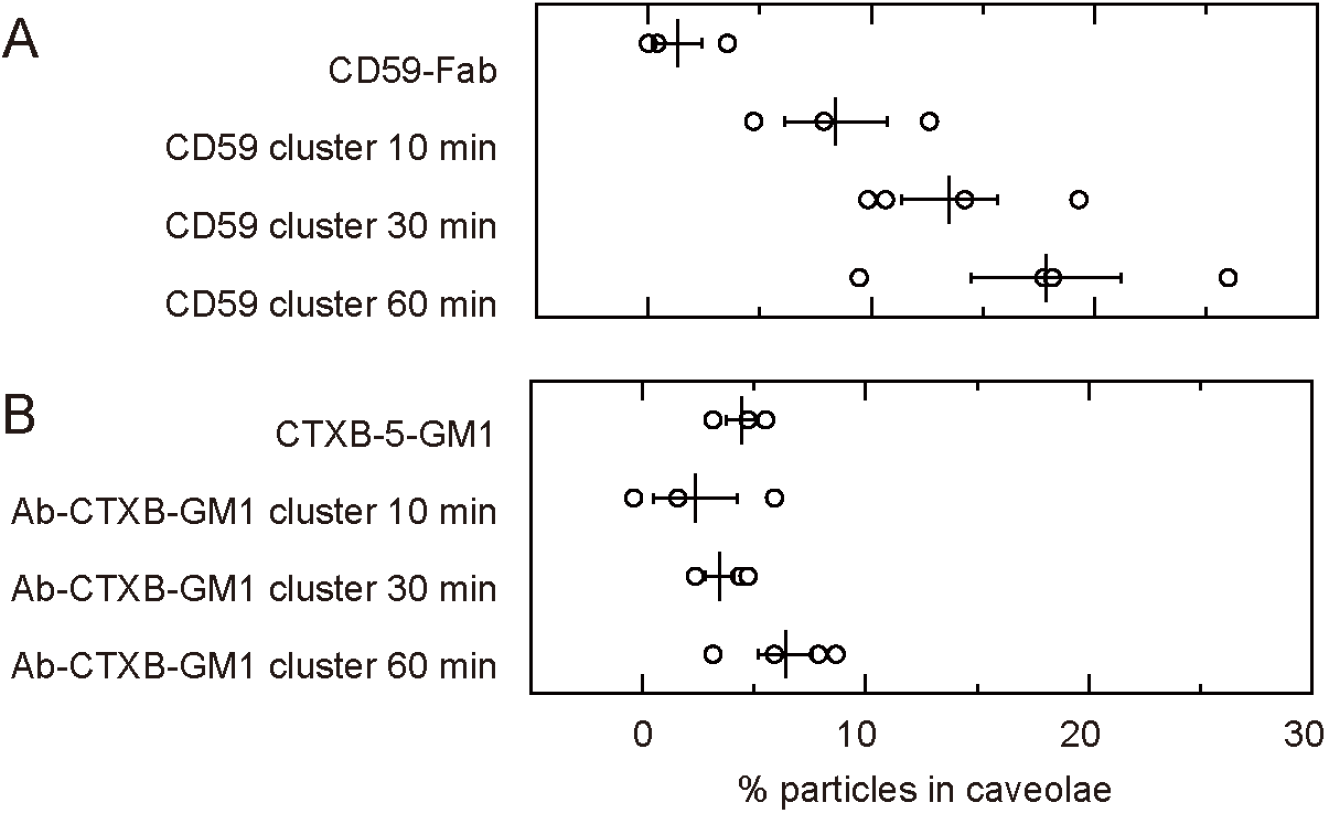
Less than 10% of CD59-cluster rafts and Ab-CTXB-GM1 clusters became trapped in caveolae within 10 min after their induction. The fractions of CD59-cluster rafts **(A)** and Ab-CTXB-GM1 clusters **(B)** colocalized with the caveolin1-GFP spots, as detected by simultaneous, two-color imaging at the single-molecule sensitivity. The colocalized fractions increased with time, but remained at less than 10% within 10 min after the cluster-formation initiation. Based on these results, all of the experiments for observing the recruitment of intracellular molecules at CD59-cluster rafts and Ab-CTXB-GM1 clusters were performed within 10 min after cluster induction.

**Figure S2.**
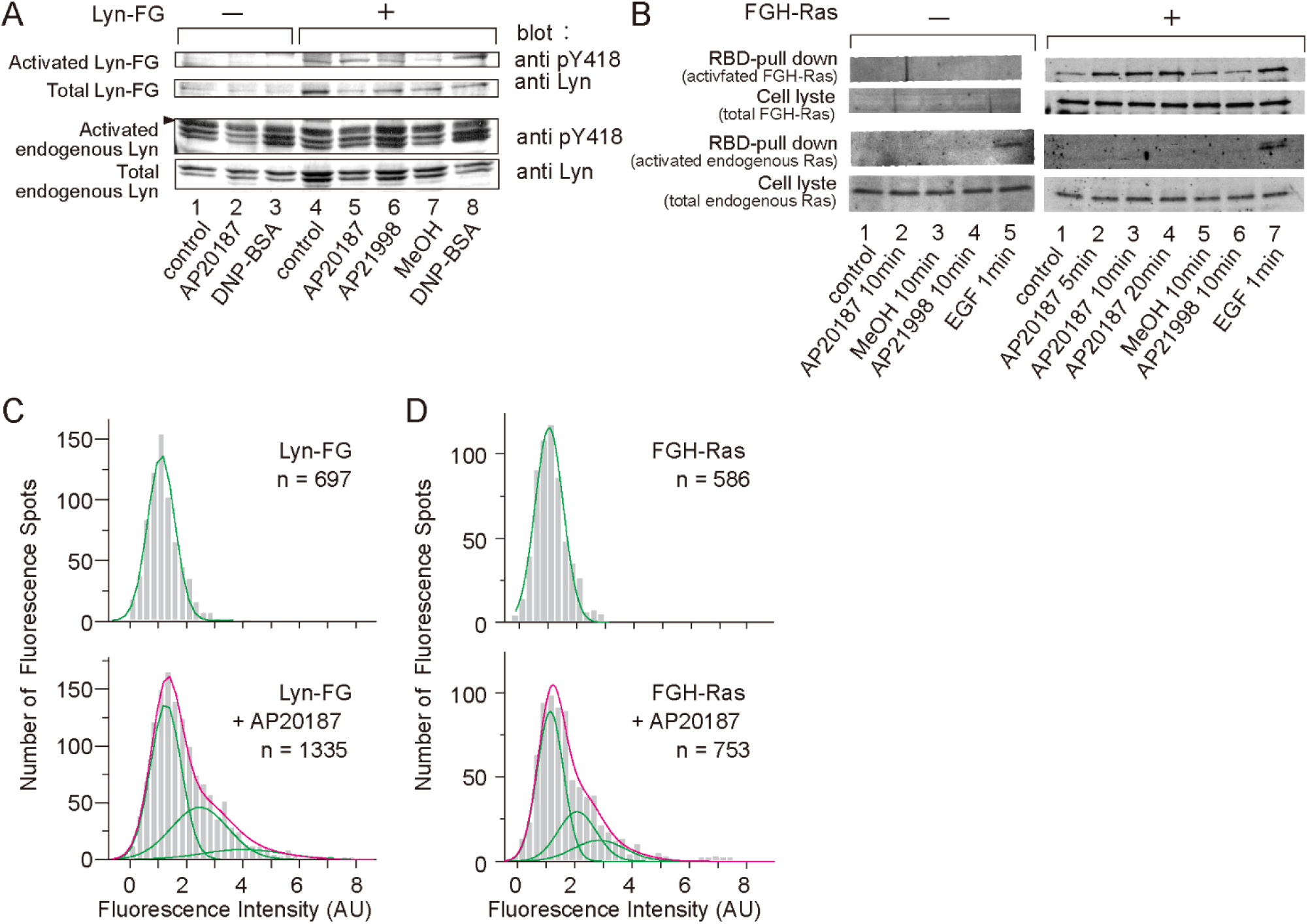
Small clusters of FGH-Ras and Lyn-FG formed in the inner-leaflet triggered signal transduction. **(A)** Lyn-FG is likely functional as it exhibited self-phosphorylation (activation) similar to endogenous Lyn in RBL-2H3 cells, after stimulation using anti-DNP IgE + DNA-BSA. Meanwhile, Lyn-FG oligomers induced by AP20187, an FKBP crosslinker, failed to activate Lyn-FG. Phosphorylation was detected by using the anti-pY418 antibodies (taking the ratio of the anti-pY418 band vs. anti-Lyn band). **(B)** FGH-Ras is likely functional as it was pulled down, like endogenous Ras, by the Ras-binding domain (RBS) of the downstream molecule c-Raf kinase bound to polystyrene beads (detection with anti-Ras antibodies), after EGF stimulation. FGH-Ras oligomerization by the addition of AP20187 induced FGH-Ras activation. AP21998 and methanol (MeOH) are negative controls. **(C, D)** Histograms showing the distributions of the signal intensities of individual fluorescent spots of Ly-FG **(C)** and FGH-Ras **(D)**, before (**top**) and after (**bottom**) the addition of AP20187. Based on these histograms, we concluded that each Lyn-FG cluster and FGH-Ras cluster contains an average of ~3 Lyn-FG and FGH-Ras molecules, respectively.

**Supplementary Table 1.** Summary of the colocalization lifetimes (*τ*_1_, *τ*_2_) and statistical parameters for the recruitment of cytoplasmic lipid-anchored molecules at CD59 clusters located in the outer leaflet.

| CD59 clusters or non-crosslinked CD59 | Cytoplasmic molecules | $\tau_1^*$<br>(ms)<br>(%) | $\tau_2$<br>(ms)<br>(%) | P-values | N for correct overlay | N for shifted overlay |
| --- | --- | --- | --- | --- | --- | --- |
| Clusters | Lyn-FG | 15 ± 0.93<br>74 | 80 ± 25<br>26 | 0.00076 | 245 | 156 |
| Non-crosslinked | Lyn-FG | 19 ± 1.3<br>100 | None | 0.86 | 122 | 83 |
| Clusters | Myrpal-N20Lyn-GFP | 20 ± 1.2<br>78 | 66 ± 58<br>22 | 0.025 | 210 | 127 |
| Clusters | TM-Lyn-GFP | 17 ± 0.92<br>100 | None | 0.46 | 93 | 127 |
| Clusters | Palpal-N16GAP43-GFP | 14 ± 1.1<br>76 | 71 ± 36<br>24 | 0.0023 | 164 | 115 |
| Clusters | GFP-C5Rho-gerger | 10 ± 0.98<br>100 | None | 0.97 | 91 | 69 |
| Clusters | FGH-Ras | 27 ± 3.3<br>73 | 91 ± 80<br>27 | 0.029 | 173 | 90 |
| Non-crosslinked | FGH-Ras | 25 ± 1.3<br>100 | None | 0.52 | 126 | 83 |
| Clusters + MβCD | FGH-Ras | 18 ± 0.78<br>100 | None | 0.41 | 536 | 468 |
| Clusters | GFP-C10H-Ras-palfar (tH) | 15 ± 1.1<br>53 | 75 ± 53<br>47 | 0.072 | 123 | 83 |

\*The mean and SEM for $\tau_1$ were determined by fitting the histogram $h(\text{incidental-by-shift})$ with a single exponential decay function. The histogram for the correct overlay was fitted with the sum of two exponential decay functions, where the decay time of an exponential function was fixed at $\tau_1$ determined for $h(\text{incidental-by-shift})$ . The fractions for the two components were calculated using $\tau_1$ and three best-fit, free-fitting parameters (two pre-exponential factors and $\tau_2$ ).

**Supplementary Table 2.** Summary of the colocalization lifetimes (*τ*_1_, *τ*_2_) and statistical parameters for the recruitment of cytoplasmic lipid-anchored molecules at Ab-CTXB-GM1 clusters located in the outer leaflet, as compared with the results at CTXB-5-GM1 and DNP-DOPE clusters.

| Outer-leaflet clusters | Cytoplasmic molecules | $\tau_1^*$<br>(ms)<br>(%) | $\tau_2$<br>(ms)<br>(%) | P-values | N for correct overlay | N for shifted overlay |
| --- | --- | --- | --- | --- | --- | --- |
| Ab-CTXB-GM1 clusters | Lyn-FG | 19 ± 0.79<br>73 | 110 ± 32<br>27 | 0.013 | 408 | 240 |
| CTXB-5-GM1 | Lyn-FG | 21 ± 1.6<br>100 | None | 0.24 | 309 | 165 |
| Ab-CTXB-GM1 clusters | FGH-Ras | 20 ± 1.2<br>86 | 97 ± 44<br>14 | 0.025 | 160 | 109 |
| CTXB-5-GM1 | FGH-Ras | 34 ± 4.5<br>100 | None | 0.52 | 60 | 31 |
| Ab-CTXB-GM1 clusters<br>+ MβCD | FGH-Ras | 26 ± 3.5<br>100 | None | 0.96 | 67 | 68 |
| DNP-DOPE clusters | FGH-Ras | 21 ± 1.5<br>100 | None | 0.39 | 86 | 81 |

\*See Supplementary Table 1.

**Supplementary Table 3.** Summary of the colocalization lifetimes (*τ*_1_, *τ*_2_) and statistical parameters for the recruitment of the outer-leaflet molecules, CD59 and CTXB-5-GM1, at the artificially induced oligomers of cytoplasmic lipid-anchored signaling molecules in the inner leaflet.

| Outer-leaflet molecules | AP20187-induced clusters of cytoplasmic molecules | $\tau_1^*$<br>(ms)<br>(%) | $\tau_2$<br>(ms)<br>(%) | P-values | *N* for correct overlay | *N* for shifted overlay |
| --- | --- | --- | --- | --- | --- | --- |
| Non-crosslinked CD59 | Lyn-FG | 17 ± 1.3<br>100 | None | 0.18 | 91 | 82 |
| Non-crosslinked CD59 | FGH-Ras | 24 ± 1.1<br>100 | None | 0.90 | 139 | 143 |
| CTXB-5-GM1 | Lyn-FG | 22 ± 1.5<br>100 | None | 0.071 | 153 | 120 |
| CTXB-5-GM1 | FGH-Ras | 21 ± 1.9<br>100 | None | 1.00 | 63 | 43 |

\*See Supplementary Table 1.

**Video 1.** Typical single-molecule image sequences obtained at a 6.45-ms resolution (155 Hz), showing a transient colocalization event of a CD59 cluster (magenta spots) and a single Lyn-FG molecule (green spots). Colocalization is indicated by white arrows.

